# Independent origins of spicules reconcile palaeontological and molecular evidence of the evolutionary history of sponges

**DOI:** 10.1101/2024.06.24.600355

**Authors:** Maria Eleonora Rossi, Joseph N. Keating, Nathan James Kenny, Mattia Giacomelli, Sandra Álvarez-Carretero, Astrid Schuster, Paco Cárdenas, Sergi Taboada, Vasiliki Koutsouveli, Philip Donoghue, Ana Riesgo, Davide Pisani

## Abstract

Sponges (Porifera) play a critical role in global biogeochemical processes. However, their evolutionary timeline remains unclear hampering our understanding of Neoproterozoic paleoecology. Molecular data suggest a Tonian or Cryogenian origin, while the oldest confirmed sponge fossils, represented by siliceous spicules, date to the Ediacaran. To understand this 150-million-year discrepancy, we compiled a new phylogenomic dataset, revised fossil evidence, derived a new dated sponge phylogeny, and tested whether ancestral sponges were aspiculated or not. We infer the last common sponge ancestor to have been early Ediacaran in age and aspiculated. Siliceous spicules evolved independently at least four times, and calcareous spicules at least two times in crown sponges. We performed a diversification rate analysis showing that siliceous spicules did not drive species radiations. Our study reconciles molecular and fossil evidence, indicating that while sponges emerged in the early Ediacaran, they are largely absent from the Neoproterozoic fossil record because they had not yet evolved spicules.

**Teaser:** Sponges emerged in the Precambrian, were soft-bodied and did not produce mineralized spicules, which emerged multiple times independently.

## Introduction

Sponges (Porifera) is an animal phylum composed of four lineages (Calcarea, Homoscleromopha, Demospongiae and Hexactinellida) sharing a bodyplan characterised by a system of pores (ostia) and channels that allow water to circulate through them. Sponges provide key ecosystem services, from habitat formation to nutrient and silica cycling, water filtering and substrate stabilisation^1^. Phylogenetically, sponges are either the sister group of all the other animals or else the second lineage to branch from the animal stem lineage^2,3,3–6^. As such, sponges are key to understanding the time of emergence and nature of the ancestral animal. However, early sponge evolution is the subject of controversy. Molecular clock studies and fossil biomarkers have long suggested that sponges diverged in the Tonian or Cryogenian^7–11^ anticipating a deep Neoproterozoic appearance of sponge fossils. Although there are Tonian, Cryogenian and early Ediacaran fossils with claims of sponge affinity, their interpretations as such range from speculative to contentious^12,13^. For example, anastomosing pores within Tonian carbonates have been interpreted as keratose sponges^14^ but corroborative anatomical evidence is lacking and more prosaic interpretations of these structures are more likely^15^. Late Cryogenian C_30_ steranes (fossil biomarkers) interpreted as evidence for Cryogenian demosponges – that is 24-ipc and 26-mes^9^, can also be interpreted as diagenetically altered C_29_ sterols from chlorophyte algae^16^. Pores in skeletal fragments from the Cryogenian of South Australia have been interpreted as sponge-like ostia^17^ but again lack corroborating anatomical evidence, while the middle Ediacaran *Eocyathispongia qiania*, originally interpreted as an adult asconoid sponge^18^, appears to lack the expected internal structure^19,20^. Finally, arsenopyrite crystal aggregates from the latest Ediacaran of Mongolia^21^ have been misinterpreted as siliceous sponge spicules^22^. There is therefore a minimal mismatch of 150 Myr between most molecular estimates for the timing of origin of crown sponges and the first unambiguous fossil evidence of sponges, represented by disarticulated siliceous spicules from the latest Ediacaran ca. 543 Ma^12,13,23,24^.

Three sponge lineages (Hexactinellida, Homoscleromorpha and Demospongiae) make spicules out of silicic acid, and a series of previous groundbreaking molecular and genomic analyses have shown that they use different biochemical pathways, suggesting that siliceous spicules might have evolved multiple times independently^25–28^. The fourth sponge lineage, Calcarea, makes spicules using calcite^29,30^. Calcareous spicules are also found in the Demosponge *Vaceletia*, both Calcarea and *Vaceletia* use carbonic anhydrase to precipitate calcium carbonate. This enzyme is also responsible for the direct precipitation of calcium carbonate from seawater in aspiculate demosponges with calcareous skeletons (e.g. sclerosponges). However, carbonic anhydrase plays a key role in many cellular functions (including the regulation of intracellular pH) and is not specific to biocalcification^31^. Furthermore, it is homologous across all Metazoa and more broadly eukaryotes^30–32^, hence its universal distribution across Porifera is not surprising and does not constitute evidence for a single origin of biocalcification in sponges.

While the oldest fossil spicules (ca. 543 Ma; Late Ediacaran) are made of silica^12,23,24^, they only slightly predate a broader diversity of early sponges (Late Ediacaran to early Cambrian) with disparate skeletal structures. Biocalcification is likely to be at least as old as biosilicification, given the existence of a diversity of Cambrian and late Ediacaran sponges with either a calcareous but aspiculated skeleton (archaeocyathids^12,33^) or biminerallic spicules made of both calcite and silica (e.g. *Eiffelia globosa*^34^). Furthermore, not all early sponges seem to have had skeletal elements. This has been demonstrated by the recent discovery of late ediacaran sponges without skeletonised parts: *Helicolocellus cantori* (539 Ma; a putative hexactinellid)^35^ and *Arimasia germsi* from the Nama group in Namibia (543-539 Ma; an archaeocyathid)^36^. Hexactinellida and Archaeocyathida^13,33^ are both crown group sponges, providing direct evidence that Porifera had a Precambrian history, although its duration remains unknown.

Fossil evidence alone paints a confusing picture of early sponge evolution, with a limited late Ediacaran record composed of a mixture of siliceous spicules and body fossils without skeletal elements, making it difficult to rationalise whether the last common sponge ancestor was spiculated^24^, and if it was, whether it had mineralised spicules made of silica, calcium carbonate or both^12^. Finally, it has also been suggested that Ediacaran sponges might have had an organic skeleton of axial filaments (similar to some keratose sponges) but no biomineralised spicules^24^, which could have led to the emergence of a long gap in the sponge fossil record. This uncertainty leaves the problem of reconciling molecular and fossil evidence for the origin of sponges unresolved.

Here we perform phylogenetic analyses of a new dataset composed of 70 genomes and transcriptomes (12 of which were newly sequenced). We use our phylogeny to perform molecular clock analyses that integrate new interpretations of the animal fossil record and recent reassessments of key fossil-bearing formations in the rock record^19^. We test whether Cambrian and Ediacaran siliceous spicules could be remains of stem or crown sponges using ancestral state reconstruction. Our ancestral state estimations integrate biochemical, morphological and fossil evidence using cutting edge structured Markov models of character state transformation with embedded dependencies^37,38^. These models allow integration of genetic/developmental knowledge when defining putative homologies across observed morphological characters, and the potential for the existence of deep homologies, underpinned by non-homologous gene replacements in ancestral regulatory networks, in ancestral state estimation^39^.

Our results indicate that sponges originated in the early Ediacaran (ca. 608 Ma), and that the last common ancestor of all the sponges did not possess spicules. Mineralised spicules emerged relatively late in sponge evolution. Calcareous spicules emerged independently in Calcarea and in the demosponge *Vaceletia*. Siliceous spicules evolved independently at least four times: once in Homoscleromorpha, once in Hexactinellida and twice in Demospongiae. In Demospongiae, siliceous spicules emerged in the stem to the Heteroscleromorpha clade and in the Verongimorpha genus *Chondrilla*, which is nested within the clade of aspiculate Demospongiae. When fossil taxa are incorporated into analyses, other independent origins of siliceous spicules emerge. Nonetheless, the last common ancestor of all sponges is invariably reconstructed as aspiculated, and we do not find evidence for an organic skeleton of axial filaments to have predated biomineralised spicules. We performed a focal study, using diversification rate shift analyses on the extant demosponges, and found two potentially significant diversification shifts nested within the Heteroscleromorpha, both postdating the emergence of siliceous spicules. Accordingly, at least in the case of Demospongiae (the most biodiverse sponge lineage) the emergence of siliceous spicules did not seem to have played a significant role in their diversification.

Our estimates of sponge divergence, combined with our study of spicule evolution and previous genomic, biochemical, and palaeontological evidence for the evolution of siliceous spicules^12,25,27,28,35^, suggest that while sponges evolved in the Ediacaran, contradicting previous analyses calibrated using fossil biomarkers, early sponges lacked spicules and had low preservation potential. Sponges had a cryptic, but relatively short, Neoproterozoic history which *de facto* reconciles the sponge fossil record with molecular estimates of divergence times.

## Results

### Phylogenomic analyses

Initially Orthofinder v. 2.7.1^40^ retrieved 5657552 genes of which the 75% were assigned to a total of 335892 orthogroups. This set of orthogroups was filtered through Phylotreepruner^41^, to remove paralogs. We decided to retain only the orthogroups that had the 90% of species left after the filtering. These resulted in 1468 orthogroups that were later filtered using the in-house built script Parafilter, to remove hidden paralogs and extremely long branch taxa. Finally, only the genes that could identify the monophyly of sponges or of the outgroup lineages identified by monophylyl.pl from https://github.com/MaxTelford/Xenacoelomorpha2019, were retained. The final dataset was composed of 70 species (64 belonging to Porifera) with the final concatenated matrix composed of 133 genes with a total length of 103,269 AAs.

Phylogenomic analyses (see Fig. 1 and Figs. S1—S3) analyses unequivocally recovered Porifera being composed of four lineages: recover Silicea (Hexactinellida plus Demospongiae) as the sister of Calcarea plus Homoscleromorpha. We recover Calcarea, with bootstrap support (BS) 100 - posterior probability (PP) 1- Coalescent branch support (CBS) 1 and Homoscleromorpha (BS=100; PP=1; CBS=0.87) form a monophyletic group (BS=100; PP=1; CBS=0.91), sister to Silicea (BS=100; PP=1; CBS=), a monophyletic group comprising Hexactinellida and Demospongiae. Within Calcarea, we recover the monophyly of Calcinea and Calcaronea (BS=100; PP=1; CBS=1). In our topologies, the Homoscleromorpha (BS=100; PP=1; CBS=0.87) is represented by two species from the two families Oscarellidae and Plakinidae, and therefore major relationships were not addressed in detail. We recovered both Demospongiae and Hexactinellida as monophyletic, with full support (BS=100; PP =1; CBS=1). For the Hexactinellida, sampling only includes representatives from the order Lyssacinosida.

**FIGURE 1.**
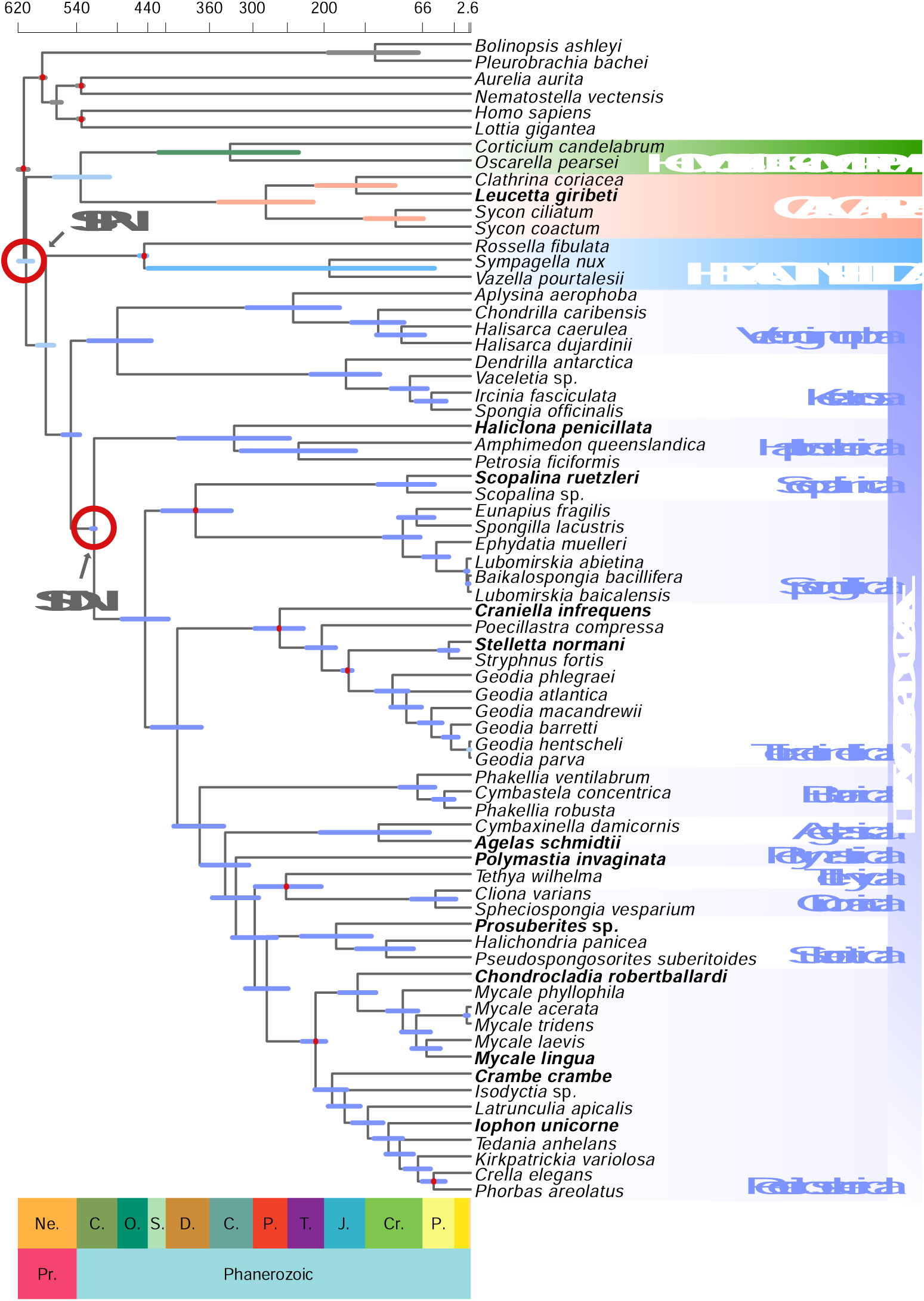
Porifera time-tree with posterior estimates using the Spicule-bearing Demospongiae (SDN) calibration strategy (red circle) and the ILN model in MCMCtree. Blue bars on nodes represent the 95% HPD. Red circles represent the two mutually exclusive calibrations strategies Spiculated Porifera Node (SPN) and SDN. Full red dots represent calibrations in both strategies. Species in bold were sequences in the framework of this study. Time scale on x-axis in hundreds of millions of years. Abbreviation on the scale bar stands for Ne, Neoproterozoic; C, Cambrian; O, Ordovician; S, Silurian; D, Devonian; C, Carboniferous; P, Permian; Tr, Triassic; J, Jurassic; K, Cretaceous; Pe, Paleogene.

In Demospongiae, Keratosa and Verongimorpha are monophyletic (BS=100; PP 0.99; CBS=0.24) and sister to the Haploscleromorpha (BS=100; PP=1; CBS=1) and Heteroscleromorpha (BS=100; PP=1; CBS=1). In the Verongimorpha, we recover *Aplysina aerophoba* (Verongiida) as the sister of Chondrillida, where we recover the genus *Halisarca* to be paraphyletic. *Halisarca dujardini* is sister (BS=100; PP=1; CBS=1) to *H. caerulea* and *Chondrilla caribensis* (BS=77; PP=1; CBS=1). Within Keratosa, we recover *Dendrilla antarctica* (Dendroceratida) to be the sister to the order Dyctioceratida, which emerges to be monophyletic and fully supported (BS=100; PP=1; CBS=1). We recover the monophyly of Haploscleromorpha (with the order Haplosclerida represented (BS=100; PP=1; CBS=1), with *Haliclona penicillata* from the family Chalinidae, being sister to *Amphimedon queenslandica* (family Niphatidae) and *Petrosia ficiformis* (family Petrosiidae), all relationships with full support (BS=100; PP=1; CBS=1). In the Heteroscleromorpha, the orders Scopalinida and Spongillida are the earliest divergent clades, forming a monophyletic clade (BS=100; PP=1; CBS=1). In Spongillida, the family Spongillidae is paraphyletic with the species *Ephydatia muelleri* that cluster sister to the family Lubomirskiidae, which is recovered monophyletic (BS=100; PP=1; CBS=1). The order Tetractinellida is recovered monophyletic (BS=100; PP=1; CBS=1). The suborder Spirophorina represented by the species *Craniella infrequens* is sister to the suborder Astrophorina. Astrophorina is monophyletic, with the family Vulcanellidae sister to the families Ancorinidae + Geodiidae. We recovered Bubarida, as sister to Agelasida, Polymastiida, Tethyida, Clionaida, Suberitida, and Poecilosclerida (BS=100; PP=1; CBS=0.96). In the BI topology, Clionaida and Tethyida formed a monophyletic sister group to Suberitida and Poecilosclerida, respectively, and the order Suberitida is recovered sister (PP=1) to the order Poecilosclerida. In the ML and CTM topology Suberitida is sister (BS=68; CBS=0.72) to the clade Clionaida + Tethyida, and Poecilosclerida is sister to these tree orders (BS=100; CBS=1). In both topologies, the poecilosclerids form two major clades, with the first one composed of the carnivorous sponge *Chondrocladia robertballardi* sister to the family Mycalidae. The second clade comprises all the other families of the order Poecilosclerida.

### Molecular divergence time analyses

Molecular divergence times (Fig. 1) were inferred for the Bayesian phylogeny (Fig. S3), using 12 fossil constraints to calibrate the ages of major sponge clades (see nodes highlighted with red circle in Fig. 1 and list for detailed descriptions of all 12 calibrations in the supplementary information). We calibrated the root to have a hard minimum of 574 Ma and a soft maximum of 609 Ma. The fossil minimum is based on the earliest occurrence of *Charnia masoni*, in the Drook Formation of Mistaken Point, Newfoundland^42^, while the maximum is based on the Lantian biota of South China which contains a diversity of macrofossils but no credible metazoans^43^. A uniform prior was used for this calibration, implying that we maintain an agnostic view of where the root of our phylogeny might lie between these constraints. The maximum on the root node is soft, with 2.5% of the probability density falling outside of the 574–609 Ma interval, on the deep side of the distribution. Eumetazoa was assigned a hard maximum of 573 Ma. This value was empirically defined to counter truncation problems in MCMCtree (see Methods for details) and a soft minimum of 561 Ma based on age of Bed B in the Bradgate Formation, where the earliest crown-eumetazoan, *Auroralumina attenboroughii*, was recovered. While it was possible to establish minimum constraints for the remaining sponge calibrations^44,45^, it was not possible to justify maximum constraints based on palaeontological, biogeographic or geological evidence. The sponge fossil record, at least in terms of records that can inform calibrations for molecular clock analyses, is based on sites of exceptional fossil preservation. As such, we have no prior expectation of whether the earliest records of various sponge clades (details in supplementary information) represent close approximations of true clade ages. Cauchy distributions with a heavy-tail (see methods) were used for these calibrations, to reflect lack of knowledge for maxima.

Two calibration strategies were used. First, we performed analyses where the oldest known siliceous spicule was used to constrain the crown Heteroscleromorpha clade. We refer to this calibration strategy as the Spiculated Demosponges Node (SDN). We then performed an analysis where the same fossil was used to calibrate the root of Porifera – Spiculated Porifera Node (SPN) strategy. These calibration strategies reflect competing interpretations of the fossil record of siliceous spicules. Under SDN, early siliceous fossil spicules are assumed to represent remains of spicule-bearing demosponges (crown Silicea), as suggested by the morphological similarity between the oldest classifiable siliceous spicules^16^ and those found in living demosponges. Under SPN, siliceous spicules are assumed to have been present in the last common ancestor (LCA) of Porifera, with the spicule fossil record providing a bound on the origin of sponges. We performed molecular clock analyses under the autocorrelated-rates (AR)^46,47^ relaxed-clock model and the independent-rates (IR)^48,49^ log-normal relaxed-clock model to assess the impact of both calibration strategies (SDN and SPN). For each analysis (i.e., SDN-AR; SDN-IR; SPN-AR; and SPN-IR), we ran six independent chains both when sampling from the prior and the posterior. We checked for convergence and used the chains that passed our MCMC diagnostics to summarise our time estimates (see Fig. S6, Tables S1-S2 and Methods for details). Under SDN, we kept three out of the 6 chains we ran under both the AR and IR relaxed-clock models (Table S1).

Results inferred using the two relaxed-clock models and different calibration strategies did not differ significantly; we therefore focus on the results obtained with the SDN-IR analysis. Results from all other analyses are presented in supplementary information (Fig. S7 to Fig. S9; Tables S3-S4). Our SDN-IR analysis inferred the last common metazoan ancestor to have emerged 616.9-605.9 Ma (early Ediacaran) (Table 1). Similarly, the sponge LCA was dated to 615.3–601.3 Ma (early Ediacaran). Crown-group Silicea (Hexactinellida + Demospongiae) was estimated at 591.3–570.3 Ma (middle Ediacaran), Lyssacinosida (within Hexactinellida) at 457.3–445 Ma (Late Ordovician), and the last common ancestor of Calcarea and Homoscleromorpha at 566.5–487.6 Ma (late Ediacaran to late Cambrian). Homoscleromorpha was estimated to have originated 429.1–228.6 Ma (middle Silurian – Late Triassic), and Calcarea 351.7–213.89 Ma (early Carboniferous – Late Triassic). The last common demosponge ancestor is estimated at 559.17–536.53 Ma (late Ediacaran – early Cambrian), and the LCA of the non-spiculated Verongimorpha and Keratosa 521.85–430.44 Ma (early Cambrian – middle Silurian). The LCA of the spicule-bearing demosponges (Heteroscleromorpha) is estimated to have existed between 521.97–515.05 Ma (middle Cambrian), and the LCA of the Haplosclerida was inferred at 403.84–246.54 Ma (early Devonian – middle Triassic). The LCA of the rest of the Heteroscleromorpha was inferred to have existed 503.17–412.61 Ma (middle Cambrian – early Devonian). Within Heteroscleromorpha, the freshwater Spongillida separated from the marine Scopalinida 424.69–325.85 Ma (late Silurian – late Carboniferous), while the heteroscleromorph lineages (Tetractinellida, Poecilosclerida and Spongillida) diverged, respectively, 299.37–231.07 (early Permian – Late Triassic), 236.53–199.74 Ma (Late Triassic – Early Jurassic) and 123.21–71.52 Ma (Cretaceous).

**Table 1.** Summary of posterior mean time estimates for all calibration strategies obtained with MCMCtree for the classes of Porifera, the Silicea clade, and Heteroscleromorpha. Full details can be found in Tables S3-S4.

| Nodes | SDN AR<br>95% CIs for time<br>estimates | SPN AR 95% CIs<br>for time estimates | SDN IR 95% CIs<br>for time estimates | SDN IR<br>95% CIs for time<br>estimates |
| --- | --- | --- | --- | --- |
| **Porifera** | 603.79-582.68 | 594.59-578.79 | 615.32-601.3 | 610.68-590.34 |
| **Silicea** | 577.48-561.72 | 567.69-551.55 | 591.32-570.3 | 575.87-542.68 |
| **Demospongiae** | 549.57-538.18 | 535.02-503.38 | 559.17-536.53 | 519.67-462.67 |
| **Hexactinellida** | 454.92-445.07 | 443.89-422.5 | 457.3-445.09 | 454.99-445.07 |
| **Calcarea** | 473.24-163.27 | 460.38-157.52 | 351.7-213.89 | 337.36-207.53 |
| **Homoscleromorpha** | 540.28-435.36 | 535.63-420.18 | 429.15-228.57 | 421.58-224.87 |
| **Heteroscleromorpha** | 521.97-515.05 | 500.69-455.96 | 520.81-515.04 | 454.95-396.53 |

### Ancestral state estimation of spicule origin

#### a. Incorporation of fossils in the phylogenomic framework

b. Ancestral character state reconstructions can be completed on a phylogenetic framework that includes only extant taxa. However, fossils can display morphological features that are not found in extant lineages, and this can impact estimates of ancestral state. Hence it is important to try to include also fossils in ancestral character state reconstructions. Unfortunately, the phylogenetic relationships of many fossil sponges is debated making their inclusion in empirical studies of the evolutionary history of spicules impossible. This is because ancestral state estimation methods and models are phylogeny-dependent, and phylogenetic uncertainty makes it impossible to place most sponge fossils in a robust phylogenetic framework. However, the recent description of *Helicocolocellus cantori*^35^ was accompanied by a formal phylogenetic analysis of morphological phylogenetic analysis that included extant and fossil taxa, providing us with a mean to include at the least some fossil sponges in our ancestral character state estimations. Importantly, these fossil sponges encompassed a relatively good diversity of early sponge spicules. These fossils are the biminerallic *Eiffelia*^50^ and *Protospongia*^51^, the siliceous *Vauxia* and *Cyathophycus*, and the aspiculated *Helicolocellus*^35^. We combined these fossils and the taxa in our phylogenomic dataset according to their phylogenetic relationships. However, in the phylogenetic tree of Wang et al.^35^, Homoscleromorpha emerges as the sister of Demospongiae, in disagreement with phylogenomic evidence^52–57^ and Fig. 1. We tested whether constraining extant taxa to follow the phylogenetic relationships supported by our phylogenomic results had an impact on the relationships of the fossil taxa. We found that when Homoscleromorpha was constrained to represent the sister of Calcarea (Fig. 1), Bayesian analyses using the same data and model of Wang et al.^35^ (see Methods) found all considered fossils to resolve as stem Silicea (Fig. 2; Fig. S10), rather than stem Hexactinellida^35^. The Cambrian siliceous spicule bearing sponge *Vauxia*, emerges as the most deeply diverging stem-silicean lineage in agreement with Wang et al.^35^. taxon in the dataset, being the first lineage to stem within total group Silicea. All the other fossil taxa emerge as member of an extinct lineage of stem-silicean. We took this instability into consideration in our ancestral character state estimations.

**FIGURE 2.**
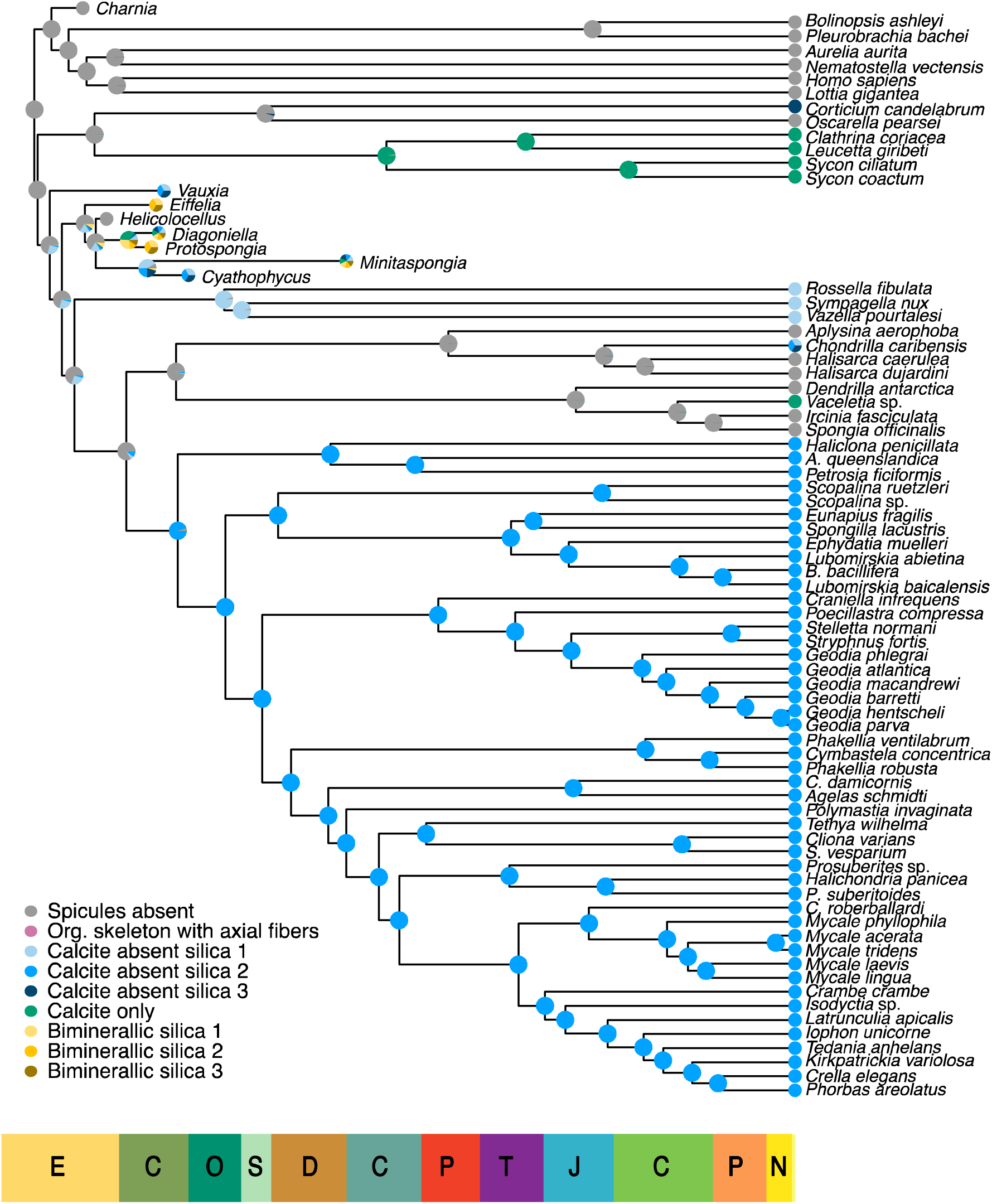
Ancestral state estimations of spicule evolution under the tree topology where fossils were allowed to optimally rearrange and the most detailed model of embedded character state dependencies (Model D). The legend depicts each colour assigned to the type of spicules mineralogy. Pink/Grey absence of spicules, blue for siliceous spicules, green for calcareous spicules, and orange for biminerallic spicules.

#### b. Ancestral state estimations

We performed ancestral state estimation using a diversity of structured Markov models with embedded dependencies^37^. These are cutting edge models of morphological character evolution, which appropriately handle the hierarchical nature of developmental and morphological character complexes^37,38^. These approaches have the potential to revolutionise ancestral character state estimation, as they allow testing whether the same morphological state, if underpinned by different developmental processes, represent convergently evolved features, or a deep homology^58^. In addition, we tested the effect of using different models of character evolution and different tree topologies (Methods for details and Fig. S11). Analyses performed under the tree topology where fossils were allowed to optimally rearrange (Figs. 2 and S10) and the most detailed model of embedded character state dependencies (Model D in Methods), strongly suggest that the last common sponge ancestor and the last common ancestor of Calcarea plus Homosleromorpha did not have spicules (P= 0.987)(Fig. 2). The crown-silicean (P= 0.64) and crown-demosponge (P= 0.768) were also most likely aspiculated (Fig. 2), even though the probability for these estimates is lower. Sensitivity analyses performed using alternative tree topologies (to account for fossil taxon instability – see above, Fig. S10 and Fig. S11) did not change our conclusions. Similarly, our results did not change if we excluded all the fossils from the analyses (Fig. S11). An aspiculated crown-sponge ancestor and an aspiculated crown-ancestor of Calcarea plus Homosleromorpha continue to have very high probability (P> 0.995), while the probability that the silicean and demosponge crown-ancestors were aspiculated increased in our sensitivity tests (P> 0.9 and P> 0.97 respectively). If we consider only extant taxa, siliceous spicules evolved independently four times (Fig. 2 and Fig. S11), once in Homoscleromorpha, once in Hexactinellida, and twice in Demospongiae (once in the stem Heteroscleromorpha and once in the verongimorph genus *Chondrilla*). If we consider also the fossils in our dataset (Fig. 2 and Fig. S11), siliceous spicules emerge independently six times, as we observe an independent origin in *Vauxia* and in the last common ancestor of *Cyathophycus* and *Minitaspongia* (Fig. 2).

Our reconstructions suggest that calcareous spicules evolved twice independently, once in Calcarea and once in the demosponge *Vaceletia*, which is nested within Keratosa. This result is interesting because our embedded dependencies models always made a strong assumption about the homology of all calcareous spicules. Yet, this hypothesis is rejected in our analyses. The inclusion of sponges with biminerallic spicules in our analyses did not influence our reconstructions of sponge ancestors deep in the sponge phylogeny, as these spicules seem to represent, at the least given the fossils in our dataset, an apomorphic condition that emerged in a clade of fossil sponges (Fig. 2, Fig. S10 and Fig. S11). Finally, our ancestral state estimations did not find any evidence that mineralised spicules might have been predated by sponges with an organic skeleton of axial filaments ^24^ (Fig. 2 and Fig. S11).

We performed a series of sensitivity tests to investigate whether coding the data using alternative structures of embedded character dependencies (Models A, B, and C in Methods), which increased the strength of our assumption of homology for the siliceous spicules, affected our results. We found that our inference of an aspiculated last common sponge ancestor, and of an aspiculated last common Calcarea plus Homosleromorpha ancestor are not affected by how we coded our data (Fig. S11). On the contrary, the condition found in the last common silicean and the last common demosponge ancestors depend on how embedded dependencies are coded (Fig. S11). These sensitivity tests imply that while our results strongly suggest that early sponges were aspiculated, our inference for the number of times when siliceous spicules independently evolved, should be considered more uncertain, as this number depends on the embedded dependencies structure imposed on the data.

### Diversification events in Silicea

We tested the effect of the origin of silicified spicules in sponge evolution, performing a case study where we inferred diversification rate shifts in Demospongiae, the most biodiverse extant sponge lineage. We followed DeBiasse et al.^59^ and first generated a phylogeny of Silicea performing a constrained ML analysis of 807 *cytochrome c oxidase 1* (COI) barcodes, using our phylogenomic tree (Fig. 1 and S3) to constrain the backbone topology (see Methods for details). The megaphylogeny was then dated (see Methods) and used to perform diversification analyses using BAMM^60^ and Medusa^61^ (Fig. S12 and Table S5; see Methods for details). BAMM identified two diversification shifts in both the AR and IR timetrees (Table S6). In the AR timetree, rate shifts are identified on the branches leading to the Axinellida plus Bubarida, and at the origin of the Poecilosclerida (Fig. 3A). In the IR topology, rate shifts were inferred on the branch leading to Hexactinellida, and again on the branch leading to the LCA of Axinellida and Bubarida (Fig. 3C). Using Medusa, diversification rate shifts were identified in the branches leading to Axinellida plus Bubarida and Poecilosclerida for both models (Fig. 3B-D), with the IR tree also identifying a shift in Dictyoceratida (Fig. 3D).

**FIGURE 3.**
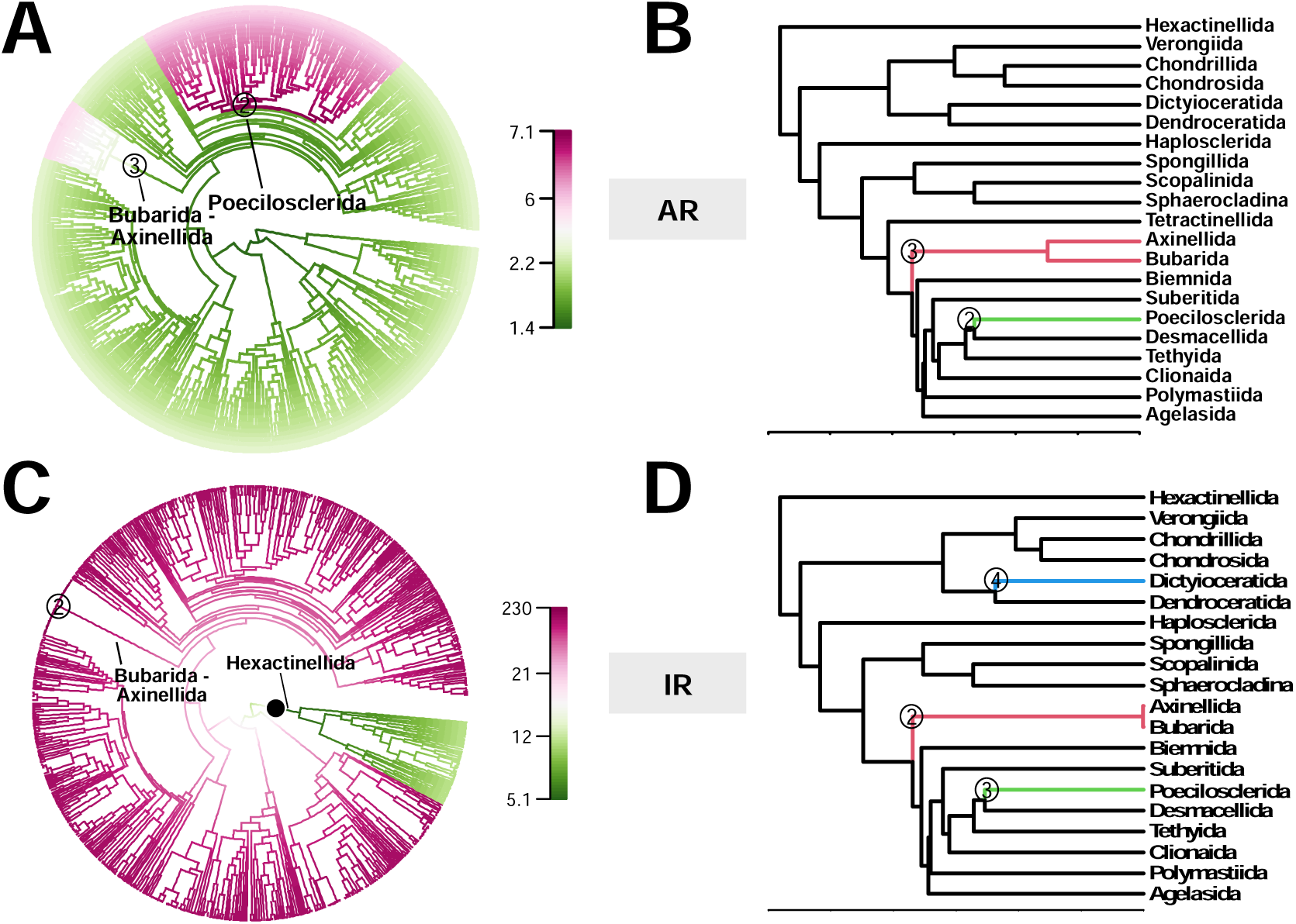
Rate diversification tests. Grey dots with numbers correspond to the shift highlighted by both Medusa and BAMM analyses, the same number is reported when the same shift corresponds on both topologies. **(A)** BAMM results for the COI timetree inferred with the AR model. The two shifts highlighted are on the branches of Bubarida + Axinellida (3), and Poecilosclerida (2). **(B)** Medusa results for the subsampled topology obtained with the AR molecular clock model. **(C)** BAMM results for the COI timetree inferred with the IR molecular clock model. The two shifts highlighted are on the branches of Bubarida + Axinellida (2) and Hexacintellida (black full dot). **(D)** Medusa results for the subsampled topology obtained with the IR molecular clock model which highlights diversification rates in Axinellida+ Bubarida(2), Poecilosclerida (3), and Dyctioceratida (4).

## Discussion

### Mineralized sponge spicules are not homologous

The results of our phylogenomic analyses confirm that sponges are split into two fundamental lineages, Calcarea plus Homoscleromorpha, and Demospongiae plus Hexactinellida (Silicea). Consistent with other phylogenomic and mitogenomic studies^53–56^, we recovered the monophyly of Keratosa, Verongimorpha and Heteroscleromorpha within Demospongiae. We resolved the aspiculate Keratosa and Verongimorpha as sister clades, and the spiculate Heteroscleromorpha as sister to Keratosa plus Verongimorpha, supporting previous findings^53^. The relationships of Bubarida have been a long-standing problem in Demospongiae phylogeny, but our tree recovers Bubarida as the sister of Agelasida+Polymastiida+Tethyida+Clionaida+Suberitida+Poecilosclerida, similar to results recently obtained with mitochondrial genomes^53^. However, we could not resolve with clear support the relationships of Suberitida with reference to Clionaida, Tethyida, and Poecilosclerida; see Figs. S1-S2.

All our timetree inference analyses recovered an Ediacaran origin of sponges and a Cambrian origin for spiculate demosponges (Heteroscleromorpha). While taxon sampling does not allow us to date the origin of Hexactinellida, our results are consistent with recent evidence for a putative crown hexactinellid (*H. cantori*) in the latest Ediacaran^35^ (539 Ma) – even though our reanalyses of the dataset compiled by Wang et al.^35^ suggests that *Helicolocellus* might be a stem-silicean rather than a stem-hexactinellid.

Our ancestral character state estimation analyses combined the taxa in our phylogenomic dataset and in the fossil dataset of Wang et al.^35^. All our ancestral state estimations reconstruct the last common poriferan ancestor and the last common calcarean plus homoscleromorph ancestor as aspiculate (P> 0.95). The condition of the silicean crown-ancestor is more uncertain, because of fossil instability (Fig. S10) and the fact that for these nodes, ancestral character state reconstructions are affected by the way in which the embedded dependencies among characters are defined (Fig. S11). Nonetheless, ancestral state analyses estimate the silicean crown ancestor as aspiculate (P= 0.66 or P=0.768) depending on what tree topology we used to represent the relationships of the fossil taxa (Fig. S11), when we code the siliceous spicules of Demospongiae, Homoscleromorpha and Hexactinellida using three different states to reflect developmental differences and biochemical differences.

Independent origin of siliceous spicules is suggested in stem-Hexactinellida, Homoscleromorpha (in stem-Plakinidae), in Demospongiae (in stem-Heteroscleromorpha) and the verongimorph *Chondrilla*. If we include fossils, two more independent origins of siliceous spicules are inferred (in *Vauxia* and in the LCA of *Cyatophycus and Minitaspongia*). This result is consistent and reinforces current interpretations of spicule evolution based on comparative genomics and biochemistry, where developmental differences are considered to represent evidence for the convergent origin of these skeletal structures^25,27,28,62^. Furthermore, our analyses do not find support for the hypothesis that early sponges might have been predated by sponges with an organic skeleton composed of axial filaments^24^. Rather, we find that stem sponges were aspiculated and that spiculated sponges evolved relatively late in sponge evolution. It is to be noted that by using structured Markov models with embedded dependencies for our ancestral state estimations, we were able to integrate biochemical and developmental knowledge in our analyses. This implies that, despite distinguishing siliceous spicules from Homoscleromorpha, Hexactinellida and Demospongiae as underpinned by different biochemistries (in our most detailed structured model – Model D in methods), we allowed for the possibility that all those spicules, and all siliceous spicules found in fossil sponges, could have been homologous. Nevertheless, our results strongly rejected that hypothesis (Fig. 2 and Sup Fig. 11).

Calcareous spicules were inferred to have emerged twice independently in our analyses, once in Calcarea and once in *Vaceletia*. This result was obtained in all our analyses even though we coded all calcareous spicules with one single state, based on the similarity of the biochemical process of biocalcification across Porifera (and Metazoa more broadly), which invariably relies on carbonic anhydrases to precipitate calcium carbonate. Rejecting the monophyly of calcareous spicules does not necessarily equate to rejecting a single origin of carbonic anhydrase-based biocalcification in Porifera. Extant and extinct sponges exist that build calcareous skeletons (e.g. the sclerotised demosponges and the archaeocyathids), but do not synthesise calcareous spicules. Current knowledge indicates that all extant sponges with a carbonate skeleton (whether spiculated or not) use carbonic anhydrase^30^, but this is not a surprising result as carbonic anhydrase has a broad distribution across Metazoa and eukaryotes more broadly, it plays a role in a variety of key physiologic functions (including pH regulation), and has been independently co-opted in biocalcification pathways multiple times across Metazoa^31^. Accordingly, while phylogenetic bracketing implies that carbonic anhydrase should have been present in the sponge crown-ancestor, the presence of this enzyme at the root of the sponge tree does not imply that the sponge crown-ancestor had calcareous spicules. In contrast, and consistently with our conclusion that calcareous spicules evolved independently twice, some genes that are involved in calcareous spicule formation (speculin, diactinin, triactinin^30^) could not be identified in our new *Vaceletia* transcriptome, although it has been noted that these proteins (which were identified in *Sycon*), seem to evolve rapidly and could not be identified also in other Calcarea^30^. Overall, we do not find any evidence to suggest that the biocalcification process in Calcarea and *Vaceletia* is underpinned by the same developmental pathway, which is inconsistent with the hypothesis that calcareous spicules in Calcarea and *Vaceletia* are homologous. These observations are consistent and support the results of our ancestral state estimation analyses. The Cambrian fossil record includes sponges bearing biminerallic spicules composed of both calcite and silica. According to our results, these spicules seem to represent a specialised condition that does not affect our reconstruction of ancestral sponges.

Given that Homoscleromorpha are always resolved as much younger than the total group Silicea in our analyses, we can only conclude that the latest Ediacaran and Cambrian spicules were most likely produced by sponges belonging to the total group Silicea, post-dating the origin of crown sponges. Given that stem- and early crown-sponges lacked siliceous spicules, they would have been unlikely to fossilize, even if they had calcitic skeletons^12^. Inevitably, our conclusions are dependent on the sampling of living and fossil sponges included in our ancestral state estimation analyses and our knowledge of their skeletal mineralogy. We recognise that there is a diversity of other fossil sponges that might be relevant to the discussion but that we could not include in our analyses, including for example the biminerallic *Lenica*, and *Takakkawia* an early sponge with a tetraradial symmetry which is unique among sponges^12^. However, we have attempted to control for this uncertainty by including a broad range of fossil taxa, controlling for the inclusion of at least a proportion of the states that are not observed in extant lineages (e.g. biminerallic sponges) but are found in fossils, including fossil taxa with siliceous spicules, and an aspiculated fossil sponge. Future research would certainly test the validity of our results, as a better understanding of the phylogeny of these sponges will emerge. Our results are consistent with the view that the different biochemical pathways used by sponges to produce siliceous spicules represent evidence that the homoscleromorph, hexactinellid and demosponge spicules are not homologous^25,27,28,63^. In detail, demosponges use silicateins enzymes^26,27,64–66^ and hexactinellids use glassin, perisilin, and hexaxilin with the organic part of the spicule (the axial filament) having different compositions^25,28,67^. Knowledge of the enzymes used by Homoscleromorpha is missing, but homoscleromorphs lack key biosilication genes found in Hexactinellida and Demospongiae, pointing to a different developmental pathway^27,68^. Furthermore, *in vitro* experiments in demosponges, particularly with *Suberites domuncula*, have demonstrated that the self-assembly of silicateins is a fundamental step in the process of demosponge spicule biogenesis itself, where scaffolds are created where silica is immediately deposited^26^. The implication is that, at least in demosponges, spicules seem to only develop as biomineralised structures.

### The emergence of silicified spicules did not drive diversification events

Sponges are globally distributed and comprise ca. 9,700 species^69^. In terms of biodiversity, Demospongiae has undoubtedly been the most successful sponge lineage^69^. Furthermore, they have been ecologically more successful, for instance being able to transition to freshwater and undergo revolutionary body plan rearrangements, leading to the only known case of carnivory in the phylum^70,71^. Many explanations have been proposed to explain their success. One is that the high diversity of spicule shapes would have been advantageous for structural functions and defence, driving adaptive radiations^72^; the results of our diversification analyses suggest otherwise. While spicules certainly play a role in defence and structural support, their emergence does not appear to have underpinned adaptive radiations, evidence of which should be shown as a diversification event on the branch that leads to the radiation of Heteroscleromorpha. Instead, we retrieve diversification events, consistently across BAMM and Medusa, on the branch that leads to Bubarida and Axinellida (Fig. 3). Both of these groups are nested well within Heteroscleromorpha, and spicules were not an innovation specific to them. Furthermore, their spicule types are shared across many other demosponge orders (Table S5) where we did not identify diversification events. Analyses that used Medusa highlighted a diversification event also in stem-Poecilosclerida in both the AR and IR timetrees. While Poecilosclerida represents the most speciose demosponge order, with ca. 2,488 species described to date^69^, with a worldwide distribution and substantial ecological diversity (evolving also carnivory – family Cladorhizidae), this result was not replicated in our BAMM analyses. In any case, Poecilosclerida are also members of the spiculate lineage and did not evolve spicules independently, although they do present one of the largest spicule complements in Porifera. We therefore conclude that while the poecilosclerid evolutionary success might have been linked to the massive expansion of microscleres observed in this clade^73,74^, the origin of spicules itself cannot have driven their diversification.

### An Ediacaran origin of sponges

Our evolutionary timescale infers crown-group sponges to be significantly younger than in previous studies (617.91–581.39 Ma early-middle Ediacaran). Our newly estimated, and younger, clade ages are driven by both a revised geochronology and reinterpretation of the fossil evidence used in establishing calibrations to be used in Bayesian node-dating analyses^19^. Previous evolutionary timelines based on molecular data have inferred sponges to have originated in the Cryogenian^8,10,11,75^. There are a number of speculative interpretations of fossil sponges from the Tonian and Cryogenian but none of these are credible^12,13^. In particular, steranes (24-ipc and 26-mes) found from the late Cryogenian (660–635 Ma) to the Early Cambrian have had a long history of alternative interpretations^7,9,76–78^, with the most recent study interpreting them as diagenetic derivatives of C_29_ sterols from chlorophyte algae^16^. Interpreting steranes biomarkers as demosponge-specific would have material implications on our evolutionary timescale. However, ambiguity over their nature implies that, unless new evidence emerges, their use to calibrate molecular clock analyses is unwarranted.

Understanding the siliceous spicule fossil record and reconciling its history against the timescale of sponge evolution is key to elucidating sponge evolution. Silica is chemically stable under low-temperature and high-pressure conditions, unlike calcium carbonate^12^. If sponges bearing siliceous spicules existed in the Neoproterozoic, they should have been preserved, given also that the Neoproterozoic ocean geochemistry was conductive to the preservation of silica, as confirmed by the presence of abundant cherts in the late Ediacaran^79,80^. The probability that sponges bearing siliceous spicules existed significantly earlier than the oldest known records (ca. 543 Ma) but were not preserved is therefore low^12,23,24^. However, the interpretation of the siliceous spicules record is difficult. The oldest siliceous spicules could be interpreted to represent evidence for spicule-bearing stem sponges, in which case the siliceous spicule record would place a limit on the age of sponges, which would be unlikely to be much older than the oldest record of fossil spicules^12^. However, these spicules can also be interpreted to represent Neoproterozoic evidence for one of the crown lineages bearing siliceous spicules: i.e. Silicea or Homoscleromorpha (although this latter is unlikely^12^, and see our time scale – Fig. 1). Under this second interpretation, the oldest spicule record would not be representative of the age of the oldest sponges. If the ancestors of these spicule-bearing crown sponges were aspiculate, they would have had low fossilisation potential, and would have been unlikely to leave a fossil record^24^. Under this second hypothesis Porifera could have enjoyed a long, cryptic, Neoproterozoic history, potentially extending to the Cryogenian or the Tonian.

Our analyses did not use the Cryogenian-to-Cambrian fossil biomarkers to calibrate our molecular clock analyses. However, our results indicate that total-group sponges have a Neoproterozoic history, albeit it only extends to the early-middle Ediacaran (617.91–581.39 Ma), that is unrepresented in the fossil record. Accordingly, we shorten the gap between the earliest unambiguous fossil evidence for sponges and molecular divergence times from ca. 150 Ma to 38–75 Ma. Our ancestral character state estimates indicate that stem sponges were aspiculate, rationalising the Neoproterozoic gap between our molecular timescale and the fossil record of early sponge evolution as a consequence of the low preservation probability of aspiculate stem-and early crown-sponges (like extant Verongimorpha and Keratosa).

In accordance with the fossil record, our results suggest that demosponge spicules emerged in ca. 31 Ma interval between 548–517 Ma (Fig. 1). In Homoscleromorpha, silicified spicules were only acquired by the Plakinidae, after the divergence from Oscarellidae in the Carboniferous. Inadequate sampling prevents us from defining precisely the period during which spicules emerged in Hexactinellida. It could be tempting to interpret the aspiculate *Helicocellus* to inform a maximum bound on the evolution of spicules in Hexactinellida. However, our reanalysis of Wang et al.^35^ indicates that this is unwarranted, as *Helicocellus* might represent a stem-Silicea rather than a stem-Hexactinellida (Fig. 2 and Fig. S10). Phylogenetic bracketing implies that hexactinellid spicules cannot be older than the silicean crown-ancestor (577.48–561.72 Ma), from which both the Hexactinellida and Silicea total-groups emerged. We conclude that stem-Hexactinellida might have been the first lineage of extant sponges to evolve spicules. However, our phylogenetic analyses (Fig. 2) confirm the siliceous-spicule bearing fossil sponge *Vauxia* to represent a stem-group Silicea, providing evidence that the oldest lineage of siliceous spicule bearing sponges were stem-Silicea (Fig. 2 and Fig. S10). Overall, our results explain the existence of an early siliceous spicule fossil record without the need to assume that stem sponges synthesised siliceous spicules.

Since sponges branch early within the animal phylogeny (irrespective of the current controversy on the relative relationships between sponges and ctenophores^2–6,55,56,81^), they are informative of early metazoan evolution, and the origin of animal multicellularity. Advocates of a literal reading of the fossil record would suggest that the origin of sponges (and Metazoa) corresponds closely with the first appearance of siliceous spicules in the fossil record (ca. 543 Ma; latest Ediacaran)^13^. Our divergence time analyses reduce by more than half the perceived gap between the earliest fossil sponge spicules and molecular estimates for the origin of sponges and this gap might diminish even further if older, unambiguous, sponge fossils are discovered. While we agree on the importance of fossils for inferring character evolution and resolving phylogenetic controversies^82,83^, our study demonstrates that the power of the fossil record in inferring evolutionary history is predicated on restoring extinct species to the rightful place within the phylogenies of living taxa. Only with an integrated understanding of phylogeny and character evolution can we hope to achieve an understanding of the evolutionary assembly of animal bodyplans^84^.

## Materials and Methods

### Sample collection, RNA extraction, library preparation, Illumina sequencing

Twelve novel sponge specimens were collected for RNA extraction (supplementary information Table S7). Samples were preserved in RNAlater (Thermo Scientific, Waltham, USA) using an overnight incubation at 4°C before long-term storage at -80 °C. Total RNA was extracted with TRIzol (Ambion, Austin, USA) and mRNA purified with a Dynabeads mRNA DIRECT kit (Thermo Scientific, Waltham, USA). The cDNA libraries were constructed with either the kits TruSeq v2 or Scriptseq v2 RNA Library Prep kit (Illumina), according to the manufacturer’s instructions, using the maximum quantity of mRNA allowed, which was 50ng – 100ng of starting material. The quality of the libraries was checked with an Agilent TapeStation 2200 system (Agilent Technologies) and the quantity with Qubit™ dsDNA HS Assay kit (ThermoFisher Scientific). The sequencing was conducted with an Illumina NextSeq 500 platform at the Sequencing Facilities of the Natural History Museum of London (Research Core Labs), using a paired-end read strategy (bp length: 2x150 bp).

### Transcriptome assembly and annotation

In addition to the 12 novel transcriptomes, 58 published RNA-seq data sets with Sequence Read Archive (SRA) IDs were included. Alla data were translated to proteins with the final total dataset composed of 70 proteomes (Table S7). For all transcriptomes assembled, read quality assessment was performed with FastQC^85^ with trimming performed with Trimmomatic^86^ with the following settings: ILLUMINACLIP:./Adapters.fa:2:30:10 LEADING:3 TRAILING:3 SLIDINGWINDOW:4:28 MINLEN:36, with the Adapters.fa file adjusted to include the adapter sequences specific to each read pair. Assembly of the clean paired reads, obtained from Trimmomatic, was performed using Trinity version^87^ with the following settings: --normalize_reads –CPU 8 –inchwormcpu 8. Prior to orthology inference, nucleotide sequences were translated using TransDecoder (https://hpc.nih.gov/apps/TransDecoder.html) for a total of 70 species and isoforms were collapsed using CD-HIT^88^. The final total dataset was composed of 70 proteomes (Table S7).

### Orthology inference

Orthologs were inferred using Orthofinder v. 2.7.1^40^, which identified 335,892 orthogroups. Putative paralogous sequences were removed using PhyloTreePruner^41^, only orthogroups with at least 90% of the species in the total dataset were retained for further analyses to minimise missing data, for a total of 1,468 orthogroups retained. Hypervariable regions of the sequences were removed using PREQUAL^89^. Orthogroups were aligned using MUSCLE^90^. Final trimming was performed with TrimAl^91^ with -auto option for each gene family. After trimming, sequences shorter than 25 AA and with less than 50% of species were removed with Al2Phylo^92^. Single-gene trees were built using IQTree v 1.6.12^93^, where a model test was run for each gene tree with the parameters -m MFP -mset LG+F+G, WAG+F+G, JTT+F+G, GTR20 -madd LG+C20+F+G, LG+C10+F+G, LG+C30+F+G, LG+C40+F+G, LG+C50+F+G, LG+C60+F+G, C10, C20, C30, C40, C50, C60, LG4M, LG4X, to allow selecting across compositionally site homogeneous and site heterogeneous models. We further used an in-house developed script (https://github.com/mgiacom/ParaFilter) by Mattia Giacomelli to remove putative hidden paralogs and extremely long branched taxa. Finally, we removed gene trees that could not identify the monophyly of sponges or of the outgroup lineages using monophylyl.pl from https://github.com/MaxTelford/Xenacoelomorpha2019. The final dataset was composed of 133 genes that were concatenated in a super-alignment using the catfasta2phyml.pl script from https://github.com/nylander/catfasta2phyml, for a total length of 103,269 AA.

### Phylogeny inference

The super-alignment was analysed using Maximum Likelihood (ML) in IQTree2 v2.1.3^94^. Model testing was performed using Model Finder plus (MFP option in IQTree but considering only across-site compositionally homogeneous models)^95^. Across-site compositionally heterogeneous models were not considered because analyses using such models were performed in Phylobayes instead (see below). Model selection identified the Q.INSECT+I+G4 model as the best fit across-site compositionally homogeneous models, we used the ultrafast bootstrap (1,000 replicates)^96^ to estimate support for the nodes in the across-site compositionally homogeneous ML trees. In addition, we ran Phylobayes^97^ under the CAT Poisson model^98^ with support values estimated using Posterior Probabilities (PP). We ran two independent chains for over 7,000 generations (Burnin = 1500). Convergence was tested using *Bpcomp* (maxdiff = 0.0018315) and *Tracecomp* (minimal ESS 61 and maximum real_diff=224488) which are part of Phylobayes (See details in table S8). Finally, we used ASTRAL^99^ as a coalescent tree reconstruction method of the final 133 gene trees, with default parameters.

### Timetree inference

After inferring individual gene trees for each available gene alignment (see orthology inference section), we ran ModelFinder to find the corresponding best-fitting evolutionary model. Following the results of model selection, we generated a five-partition sequence alignment (see supplementary information for justifications). Prior to timetree inference, we used CODEML (part of the PAML package^100^) to estimate the vector of branch lengths, the gradient (i.e., vector of first derivatives of the likelihood function), and the Hessian (i.e., matrix of second derivatives of the likelihood function) for each partition under maximum likelihood. These vectors and matrices are necessary to subsequently enable the approximate likelihood calculation in MCMCtree^100^ (see supplementary information for details).

We fixed the tree topology to our inferred Bayesian tree (Fig. 1 and Figs. S3, S12) and constrained the age of 12 nodes representing major sponge clades in our phylogeny (red dots and circles in Fig. 1) based on fossil evidence. The oldest fossil spicule was initially used to calibrate the origin of the Spicule-bearing Demospongiae (SDN calibration strategy). We used a uniform prior on the root age with a soft maximum of 609 Ma (pU = 0.025) and a hard minimum of 574 Ma (pL = 1e-300; this small value enables the usage of hard bounds in MCMCtree as pL=0 would cause numerical underflow). We constrained the age of Eumetazoa with a hard maximum of 573 Ma (pU = 1e-300) and a soft minimum of 561 Ma (pU = 0.025) to avoid truncation issues during the MCMC runs^75^. The age of the other ten calibrated nodes was constrained using heavy-tail Cauchy distributions (i.e., lower-bound calibrations with an offset of p=0.1 or p=0.5, a scale parameter of c=0.5 or c=0.1, and a left tail probability of pL=1e-300 to enforce a hard bound on the specified minimum age). The use of a heavy-tailed distribution was preferred as it allows us to capture the credibility of the sponge fossil record, which is unlikely to be reliable in deep time because of taphonomic processes having altered diagnostic characters, thus posing further difficulties on taxonomic classifications. We performed a sensitivity test (SPN strategy) where the oldest siliceous spicule was reassigned to calibrate the node representing the last common sponge ancestor (see also Fig. 1) to account for the uncertainty in the spicule fossil record, allowing us to investigate its impact on poriferan divergence times derived from a molecular clock-dating analysis. In short, the SDN strategy constrained the spicule bearing Demospongiae (Heteroscleromoprha) to have a hard minimum at 515 Ma, while the SPN strategy constrained the root of Porifera to a minimum age of 515 Ma. Both experiments used a Cauchy distribution specified in the same way (c= 0.5, p = 0.5, and pL=1e-300). The age of 515 Ma used in both the SDN and SPN strategies is based on loose siliceous spicules from the lower Cambrian (Series 2, Stage 3) Sirius Passet Biota of North Greenland^101^. A list of all the calibrations used can be found in the supplementary information.

We ran all analyses under both the IR^47^ and the AR^46,47^ relaxed-clock models implemented in MCMCtree (PAML package v4.9i^100^). Six independent chains for every analysis (SDN+AR, SDN+IR, SPN+AR, SPN+IR) were run with 4,020,000 iterations each. As per our MCMC settings, we discarded the first 100,000 collected samples as part of the burn-in phase and, every 100 iterations, we saved the values sampled for each parameter until reaching a total of 20,000 samples for each chain. The MCMC settings for the evolutionary model can be found in the supplementary information. We run six MCMC chains for both our prior and posterior estimations and used an in-house pipeline (see https://github.com/MEleonoraRossi/spicules-dating) to (i) filter these chains and (ii) generate convergence plots and summary statistics. The plots and summary statistica only used the chains passing our quality control checks (see supplementary information and GitHub repository for methodological details).

### Ancestral Character State Reconstructions

#### Assembling a phylogenetic framework with fossils and extant taxa

Sponges have a rich fossil record^12^, which can provide additional evidence of spicule evolution. Fossil observations are particularly useful for ancestral state estimation, as their closer proximity to the root of the phylogenetic tree compared to living taxa gives them greater influence on ancestral likelihoods. Unfortunately, the majority of fossil sponges have never been included in a formal phylogenetic analysis and their relationships are unknown. However, the recent study of the relationships of *H. cantori*^35^, included a good selection of fossil sponges. Crucially, this study included fossil taxa with a broad diversity of skeletal structure, from the aspiculate *H. cantori* itself to the biminerallic *Eiffelia* and *Protospongia,* and the siliceous *Vauxia*. In order to test the effect of fossils on our ancestral states analysis, we produced time trees integrating the results of our phylogenomic analysis with those of Wang et al.^35^. To determine fossil placements within our phylogenomic framework, we reanalyzed the Wang et al. morphological dataset using MrBayes 3.7^102^, constraining the topology so that living taxa align with our phylogenomic results. That is, we constrained Calcarea and Homoscleromorpha to be sister classes, and likewise for Hexactinellida and Demospongiae. We then performed two constrained Bayesian analyses. In the first the fossils were left fully unconstrained so that they could arrange in their optimal position in the context of the constraints provided for by extant taxa. In the second, we constrained the fossils to be members of the same clades where they were found by Wang et al.^35^. The input files used to run these analyses are available at https://github.com/MEleonoraRossi/ASE_spicules/. This allowed us to account for instability in the relationships of the fossil taxa in our ancestral state reconstructions. Topologies were subsequently time-scaled using the timeplaeophy() function in the paleotree R package^103^, specifying the ‘equal’ method. Fossil tip ages were taken from the Wang et al.^35^ and internal node ages were fixed based on our molecular clock analyses. We generated 12 alternate timescaled trees, representing all possible combinations of three variables: (1) tree topology (a - a tree without fossils; b - fossils unconstrained; c - fossils constrained to follow Wang et al.^35^); (2) molecular timescale (a - autocorrelated rates, 2 - independent rates); (3) calibration strategy (a - SDN; b - SPN).

#### B) Estimation of ancestral states

We conducted marginal likelihood ASE in R using the fitmk() and ancr() functions from the phytools package^104^. Character states for the full dataset were retrieved from the literature (Table S9). We used Embedded Dependency Models^38^, and accounted for biochemical/developmental differences and similarities when investigating the evolution of different types of spicules. We coded the following characters: (i) Spicule: absent= 0, present=1. When a spicule was present the following conditions were possible: (ii) Spicule has calcitic mineralisation: absent =0, present =1; (iii) Spicule has siliceous mineralisation: absent =0, present =1. Finally when siliceous spicules were present, we treated their developmental pathways as alternative states: (iv) Siliceous spicule mineralisation using the pathway of Demospongiae= 1; Hexactinellida= 2; Homoscleromorpha= 3. Note that for calcareous spicules we did not distinguish different developmental pathways, implying that all sponges with calcareous spicules define their spicules using the same developmental pathway. This reflects the universal use of carbonic anhydrase by sponges to synthesise carbonaceous skeletal elements (not only spicules). We tested four ED models where characters were amalgamated in different ways: Model A = char(iI) + char(iii); Model B = char(i) + char(ii) + char(iii); Model C = char(ii) + char(iii) + char(iv); Model D = char(i) + char(ii) + char(iii) + char(iv). We accounted for transition rate heterogenity by using both Equal Rate (ER) and an All Rates Different (ARD) model. We iterated through each combination of models when amalgamating them into ED models. For example, for Model A = Char(ii) + Char (iii), we used ER and ARD on each character, so for Model A we ended up doing analyses using: Char(ii)ER & Char(iii)ER; Char(ii)ER & Char(iii)ARD; Char(ii)ARD and Char(iii)ER; and Char(ii)ARD & Char(iii)ARD. The number of embedded dependencies model combinations therefore equal 2 to the power of the number of characters being amalgamated. Each ED model was applied to each of the 12 timescaled trees (see previous section). We calculated the Bayesian Information Criterion (BIC) and Model Weight for each ED plus timescaled tree combination and plotted the model average per tree topology. Overall we performed 432 ancestral state estimations (all scripts used to perform these analyses are available https://github.com/MEleonoraRossi/ASE_spicules/).

### Diversification Rates through Time and Across Clades

We assembled a dataset of 804 COI sequences from GenBank. The sequences were visualised using Seaview and aligned with MUSCLE v. 3.8.31^90^. The final alignment was 609 bp. We inferred the best-scoring maximum-likelihood (ML) tree with IQTree2 v2.1.3^94^. However, we constrained the analyses using our phylogenomic tree (Fig. 1) as a backbone constraint^59^. Model testing was performed using Model Finder plus (MFP option in IQTree but considering only across-site compositionally homogeneous models) because of the size of the dataset which limited our ability to use complex models. Following the procedure detailed in Álvarez-Carretero et al., 2022^105^, we fitted skew-t (ST) distributions to the posterior time densities inferred using PAML v4.9i^100^ (see supplementary information). We selected 13 ST distributions that matched the nodes that had been calibrated prior to timetree inference so that we could use them as prior distributions in the subsequent analyses (Table S10) on the ML topology inferred using the 804 COI sequences. We ran 64 independent chains with 4,020,000 iterations each. We discarded the first 20,000 iterations as part of the burn-in phase and subsequently collected samples every 20 iterations. We saved the values sampled for each parameter until reaching a total of 200,000 samples for each chain. Convergence was calculated as explained in the “Timetree inference” section above (see also Table S5).

Macroevolutionary dynamics of diversification were modelled with the software Bayesian Analysis of Macroevolutionary Mixtures (BAMM) v.2.5.0^60^ on the timetree inferred following the procedure described above. We accounted for incomplete taxon sampling by defining a sampling fraction of 5%, which corresponds to the number of species included in the COI dataset compared to the total number of Silicea species accepted. For each order we also calculated the sampling frequencies (species sampled/species accepted for the order; data obtained from the World Porifera Database^69^; Table S11). Ten million generations of reversible jump Markov Chain Monte Carlo sampling were run, drawing samples from the posterior every 10,000 generations. For the prior probability of rate shift, we tested values ranging from 0.1 to 50 and chose the value leading to the highest ESS values for LogLikelihood and NumberOfShifts (Table S4). We processed the output data with BAMMtools ^106^ and, after removing the samples collected during the burn-in process (10% of the samples), we obtained the summary statistics, and plotted the diversification rate over time. To overcome the possible bias introduced by incomplete sampling in diversification analyses, we confirmed our results using the taxonomic method implemented in MEDUSA^61^ where clades are collapsed to terminal lineages of equal ranks. Only diversification events inferred by both methods were deemed reliable.

## Author contributions

DP, AR and PJD conceived the study. MER performed the analyses and wrote the first draft of the paper. DP, PJD and AR edited the paper. MG, NK, and MER performed orthology inference. JNK provided support for the ASE analyses, SAC provided support and scripts for timetree inference analyses. PC and AS provided some of the fossil calibrations and helped assemble the morphological matrix used in the ASE. ST, VK, AR, and PC took part in the sampling. BP, ST, VK, NK and MER extracted the RNA and performed library preparations of the samples.

## Data and materials availability

All data, code, and materials used in the analyses must be available in some form to any researcher for purposes of reproducing or extending the analyses. Include a note explaining any restrictions on materials, such as materials transfer agreements (MTAs). Include accession numbers to any data relevant to the paper and deposited in a public database; include a brief description of the dataset or model with the number. The DMA statement should include the following: “All data are available in the main text or the supplementary materials.”

