## Supplementary material for "Independent origins of spicules reconcile palaeontological and molecular evidence of the evolutionary history of sponges"

Maria Eleonora Rossi *et al.*

**This PDF file includes:**

Supplementary Text

Figs. S1 to S12

Tables S1 to S11

Data S1 t

Supplementary Text

**Material and Methods**

**Timetree inference analysis**

To estimate species divergence times, we dated our inferred Bayesian phylogeny (Fig. S3) using MCMCtree, which is part of the PAML package^1^, with the approximate likelihood approach^2^. CODEML, also part PAML^1^, was first used to calculate the first and second derivatives of the likelihood function (i.e., the gradient and the Hessian) and the branch lengths under maximum likelihood, which are then used by MCMCtree to approximate the likelihood function using the Taylor expansion (see PAML documentation).

To improve precision in divergence time estimation, it is best to partition a superalignment into various alignment blocks, so that each block (or partition) contains subsets of genes that share the same evolutionary process (and thus evolve at the same, or similar, evolutionary rate)^3–5^. However, the number of chosen partitions cannot be arbitrarily large; including more than 10 partitions can artificially inflate the precision with which divergence times are estimated^6^. Consequently, we ran IQtree to find the best-fitting model for each gene alignment that was part of the superalignment, but considered only those models available in PAML. A total of 5 different models were selected as the best-fitting evolutioanry models, which allowed us to group our gene alignments into five different alignment blocks^6^. For each alignment block, we selected the relevant rate matrix according to the best-fitting evolutionary model so that CODEML could estimate the branch lengths, the gradient, and the Hessian later required by MCMCtree (see PAML documentation). The first (68 taxa and 25,620 AA) and second (68 taxa and 25,620 AA) partitions used independently parameterised JTT+G4 models. These gene alignments were separately classified as IQtree identified the best-fitting models as having very different alpha parameters for the Gamma distribution that accounts for rate heterogeneity. Please note that IQtree inferred the best-fitting model of one of these alignment blocks as JTT+G4+I, which is not implemented in PAML, hence we had to simplify this model to JTT+G4 instead when using CODEML. The model for the third partition was DCMut+G4 (66 taxa and 7,388 AA). The fourth (69 taxa and 9,947 AA) and fifth (70 taxa and 49,102 AA) partitions used independently parameterised LG+G4 models; same scenario that occurred with the JTT models as aforementioned.. The main output of CODEML is a file called “rst2”, which contains the vectors and the matrix. The content of the five “rst2” files (i.e., one per partition) were then concatenated in the so-called “in.BV” file (see PAML documentation).

To estimate divergence times, MCMCtree was then run with the 5-partition in.BV file, the 5-partition AA alignment, and the fixed tree topology of Fig. S3 with 12 calibrations (see below Supplementary List of Calibrations). The prior on divergence times consists of a birth-death process with species sampling^7^ that we defined with λ = μ = 1 (birth and death rates, respectively) and a sampling fraction of ρ = 0.1. The resulting kernel density is an approximate uniform distribution that is used to determine the distribution of the node ages for uncalibrated nodes. We used a gamma-Dirichlet rate prior on the mean i-*th* partition rate, μi ~ Γ(2,20) so that the mean evolutionary rate is 0.1 substitutions per site per 100 Myr, the time unit we defined. We used a more diffuse gamma-Dirichlet prior on the variance of the rate for each locus *i*, σi2 ~Γ(1, 10), to account for the violation of the clock that is expected in deep phylogenies. For each combination of clock models and calibration strategy (SDN+AR, SDN+IR, SPN+AR, and SPN+IR – see main for abbreviations), we ran 6 independent chains. We discarded the first 100,000 iterations as part of the burn-in phase and collected a total of 20,000 samples every 100 iterations. In total, all chains ran for 2,100,000 iterations.

**MCMC diagnostics**

We assessed the quality of the six chains we ran for each analysis. We calculated the mean lower- and upper-quantiles (i.e., 2.5%- and 97.5%-quantiles) of each time distribution (i.e., mean lower- and upper-quantiles for each node in the phylogeny) for each node in Fog. S3, and each of the six chains we ran per analysis. In an iterative manner, we used our in-house R function check_quantiles to compute, for each node, the difference between the quantiles computed for each chain. This is an iterative process where we start denominating Chain 1 as the “main” chain, and calculate differences between the main chain (Chain 1 in the first instance) and the other 5 chains.   After that, the procedure is repeated while considering each one of the other six chains as the main chain.   For each node, the resulting differences are compared to a predefined threshold value, α, to establish whether the chain being compared to the “main” chain should be flagged as potentially “problematic”.  If the difference between quantiles is either too large or too low, there may have been convergence issues, and the compared chain is flagged as potentially problematic.  Chains labelled as “problematic” are not used for summary statistics nor to compute the mean divergence times. Note that whether a chain is deemed problematic or not may depend on what chain is used as “main”.  If the chain used as main is problematic, all (or almost all) of the other chains would be flagged as problematic.  Accordingly, to avoid using a potentially problematic chain as main, once all 6 chains are tested, the one that minimises the identification of problematic chains when used as the “main” chain is selected as the “reference” chain. Chains identified as problematic when using the reference chain were not used in our estimation of divergence times. Note that while for shallow phylogenies we can use a stringent threshold value (e.g., α = 0.05 or α = 0.1), deeper phylogenies require more relaxed values as the uncertainty in time estimates increases (i.e., larger differences between quantiles are expected). For our phylogeny, we used a threshold value of α = 0.25, and did not keep chains for which, for at least one node, quantile differences were either larger or lower than α = 0.25 (see supplementary tables S3-4 for the results with SDN+AR, SDN+IR, SPN+AR, and SPN+IR).

We wrote a wrapper function around the R function rstan::monitor v.2.21.7 to (i) calculate the effective sample size (ESS) for bulk and tail quantiles as well as (ii) the potential scale reduction factor on rank normalised split chains (i.e., the Rhat value) for all divergence times estimated with the samples collected by the chains that passed our filters. Chain convergence is theoretically assumed if ESS values are over 100 and Rhat values are smaller than or equal to 1.05. After summarising our parameter estimates with the filtered chains, the bulk- and tail-ESS values were over 100 and the Rhat values were smaller than 1.05 for all our model parameters of interest (see supplementary Tables S1-S2). To show the quality of our statistical QCs, Figure S6 is used to compare the convergence plots generated with all the chains and then with only those chains we kept after applying our filters. To build the plots, if 4 chains had passed our filters, the average of all the divergence times was calculated with the first 2 chains and with the last 2 chains. The estimated mean divergence times were then plotted against each other. If the resulting convergence plot shows an almost straight line (i.e., x =~ y), the chains are assumed to have converged. In the plots with unfiltered chains, some points deviate from the “x =~ y” line, suggesting potential convergence problems for some chains.  These issues disappear when only chains retained after filtering are used. All the data, scripts, and step-by-step guidelines required to reproduce these analyses can be found in our GitHub repository: https://github.com/MEleonoraRossi/spicules-dating

**Sensitivity test for fossil calibrations**

There is some disagreement on the nature of *Geoditesia jordaniensis*^8^ (Cardenas 2020).  Accordingly, we performed a sensitivity analysis where we changed the calibration interval defined by this fossil (166.1–163.5 Ma), to 166.1–150 Ma, to take into consideration uncertainty on this part of the sponge fossil record.  This analysis did not affect our results.

**Ancestral State Estimation**

Fossils might include states that are not observed across extant taxa.  This can have an impact on ancestral state estimation. Accordingly, we included as many fossils as possible in our ASE analyses.  Many fossil sponges are known, but only a few have been included in phylogenetic analyses, and including in ASE fossils for which phylogenetic relationships are unknown would be computationally too intensive as one should integrate across all possible placements of each possible fossil.   Recently, Wang et al.^9^ published an analysis of sponge fossils including a diversity of taxa with interesting skeletal characters.  The data set included biminerallic sponges as well stem sponges with siliceous spicules and one stem aspiculate sponge. We tested the effect of including fossils in our ASE using the dataset of Wang et al.^9^.  We however noted that the phylogeny published by Wang et al.^9^ disagreed with knowledge of sponge relationships with reference to extant taxa.  This is because Wang et al.^9^ tree placed the Homosceromorpha as the sister of Demospongia (based on morphological evidence), rather than as sister of Calcarea (see Fig. 1).  This result suggests that Wang et al.^9^ phylogeny might be partially driven by homoplasy, probably relating to the existence of silicified spicules in Homoscleromorpha. To account for the possible effects of homoplasy in the results of Wang et al.^9^ we repeated an analysis of their dataset (using the exact same method and model), but containing the extant sponges in their dataset (a sampling of demosponges, homoscleromorphs, hexactinellids and calcareans) to follow the relationship suggested by phylogenomic data (i.e. homoscleromorphs sister of calcareans, and demosponges sister of hexactinellids).  This was achieved using a backbone constraint in MrBayes^10^.  Fossils were left free to arrange themselves based on the data.  All other parameters in the analyses were retained exactly as in Wang et al.^9^.  We then performed our ASE on three different topologies: (1) our original phylogenomic tree (with no fossils);  (2) a tree where the extant taxa were arranged as in Wang et al.^9^ and the extant taxa as in Fig. (1); and (3) the tree that emerged from the constrained analysis described above. The rest of the methods for the ASE are reported in the main paper.

**Supplementary list of calibrations**

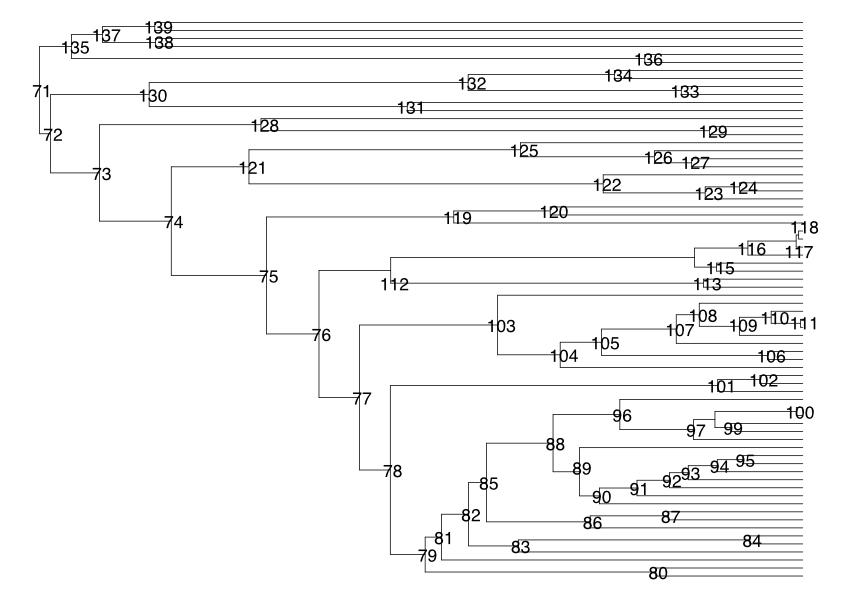

**Node 71: Crown Metazoa** | 574-609 Ma

Fossil taxon and specimen: *Charnia masoni* (LEIUG 2328 from Bed B of North Quarry, Charnwood Forest, UK).

Phylogenetic justification: Dunn et al. {, 2021 #17959}^11^ demonstrated that *Charnia masoni* is a stem-eumetazoan based on analysis of its morphogenesis and comparative taphonomy.

Minimum age justification: Matthews et al. {, 2020 #16515}^12^ date the earliest occurrence of *Charnia masoni* to the Drook Formation of Mistaken Point, Newfoundland, which has been dated to 574.17 Ma ± 0.66 Myr, providing for a minimum constraint of 573.51 Ma. In this case the age we set the minimum is 574 to avoid truncation issues as explained in the Supplementary method section.

Soft maximum age justification: We establish the minimum constraint on the Lantian biota of South China which contains a diversity of macrofossils but nothing that can definitively be classified as a metazoan. Yang et al. {, 2022 #19193}^13^ derive a Re-Os age for the Lantian of 602 Ma ± 7 Ma, resulting in a maximum constraint of 609 Ma. This age allows for the possibility of *Eoandromeda* being a crown metazoan.

**Node 135: Crown Eumetazoa** | 561.1-590.8 Ma

Fossil taxon and specimen: *Auroralumina attenboroughii* (GSM 106119) from Bed B, Bradgate Formation, Charnian Supergroup of North Quarry, Charnwood Forest, UK {Dunn, 2022 #18823}^14^.

Phylogenetic justification: Dunn et al. establish a crown-cnidarian affinity for *Auroralumina attenboroughii* based on a formal character analysis.

Minimum age justification: The age of Bed B within the Bradgate Formation has been constrained to 563 Ma ± 1.9 Myr^15^. Noble et al.^16^ later derived a U-Pb date of 556.6 ± 6.4, but the associated uncertainty entirely encompasses the original, more precise date. Hence, we follow the earlier date from Wilby et al.^15^.

Soft maximum age justification: We establish the minimum constraint on the Weng’an biota of South China which contains a diversity of microfossils, many of which have been interpreted as metazoans in the past^17^, but which have subsequently been reinterpreted as stem-metazoans, at best^18,19^. Yang et al.^20^ establish a maximum age of 587.2 Ma ±3.6 Myr for the Weng’an Biota based on the Re-Os system. This provides for a maximum constraint of 590.8 Ma.

**Node 138: Crown Cnidaria** | 531.80 Ma.

Fossil Taxon specimen: *Olivooides multisulcatus* (Geological Museum of Peking University: GMPKU3083-GMPKU3090), Dengying Formation, Fortunian Stage, constituting a range of embryonic and post-embryonic developmental stages. This calibration largely follows Benton et al.^21^, from which most of the description is derived.

Phylogenetic justification: *Olivooides* is known from embryonic and post-embryonic stages of development, including a polyp theca, characteristic of scyphozoans, and a medusa stage^22^.

Minimum age justification: *Olivooides multisulcatus* co-occurs with *Anabarites trisulcatus*, which is indicative of the middle of the Fortunian Stage of the Terreneuvian Series, the first of the Cambrian. We derive a numeric age of 531.80 Ma following the age model of Bowyer et al^23^.

**Node 139: Crown Bilateria** | 532 Ma.

Fossil taxon and specimen: *Aldanella janjiahensis* (YXII102-02), often synonymized with *Aldanella attleborensis,* from the Lower Cambrian Dahai Member of Zhujiaqing Formation, Xiaotan, Yongshan County, Yunnan^24^.

Phylogenetic justification: *Aldanella* is a dextrally coiled mollusc assigned to the Pelagiellida. The distinct asymmetries and the preservation of muscle scars^25^ suggest that it is a partially coiled stem-group gastropod.

Minimum age justification: The fossil occurrence falls fully within the *Anabarites trisulcatus–Protohetzina anabarica* Assemblage Biozone^26^, falling within the span of 537 – 532 Ma.

**Node 72 and Node 75: Crown Demospongiae** | 515 Ma.

Fossil Taxon Specimen: Demosponge indent (MGUH 30886 from GGU sample 340103.3684), from the Sirius Passet Lagerstätte, Buen Formation, Peary Land, North Greenland^27^.

Phylogenetic justification: The combination of spicule forms (monaxons with small sigma, toxa and unique spiral morphologies) places the sponge as the first crown-group demosponge, thus representing the stem lineage of Heteroscleromorpha.

Minimum age justification: The fossil occurrence falls within the Cambrian Series 2, Stage 3 and therefore presents a reliable calibration point for the minimum root age of the group Haploscleromorpha + Heteroscleromorpha which is ~ 515 Ma^27^.

**Node 88: Poecilosclerida** | 199 Ma

Fossil Taxon Specimen: Poecilosclerida, Lower Liassic Kirchsteinkalk (Allgäu Formation as basin facies links to Liassic Kirchsein Limestones) of the Northern Calcareous Alps in Germany).

Phylogenetic justification: Based on the morphological characterization of a dense occurrence of C-shaped “sigma microscleres” (type forceps) that are distinct to other known sigmas within demosponges, thus most similar to recent poecilosclerid sigmas, as well as the presence of various chelae spicules^28^. Chelae are a synapomorphy of the Poecilosclerida.

Minimum age justification: Isochelae, are for the first time undoubtfully reported from the Early Jurassic Hettangian/Sinemurian boundary 199.3 Ma.

**Node 83: Tethya** |37.8 Ma

Fossil Taxon Specimen: *Tethyastra* sp. ZPAL Pf.26, St. Vincent Basin (Blanche Point section)

Phylogenetic justification: numerous oxyasters (Figs. 13F–H)^29^ resembling those of recent *Tethyastra oxyaster* Burton, 1934. The great morphological resemblance and the occurrence of recent *T. oxyaster* from all over Australia^30^ confirm the assignment to the family Tethyidae and the Order Tethyida.

Minimum age justification: Fossils were sampled from sediments that were collected from the outcrop that is part of the mid-Eocene to mid-Oligocene, 200 m thick succession that is overlayed by Pliocene and Pleistocene sediments^31^. More detailed fossilized spicules fall within the Priabonian age 33.9–37.71 Ma. They derived from two middle units of the Blanche Point Formation: Gull Rock member (Mb.) and Perkana Member.

**Node 94 Phorbas** | 37.8 Ma

Fossil Taxon Specimen: *Crellastrina* sp. ZPAL Pf.26, Upper Eocene units sampled from Doyle Road, Princess Royal, and the Hamersley River glauconitic and spiculitic marls and limestones. Eastern South Australia.

Phylogenetic justification: Studied spicules of the fossils resemble those of recent *Crellastrina alecto* Topsent, 1898 [described as Yvesia; family Crellidae (see Fig. 22D)^29^ the most, which belongs to the family Crellidae and order Poecilosclerida^67^. Notably, it is suggested that based on the spicules it might be that there is no recent equivalent representative of this fossil, but it has been hypothesised that the spicules still resemble those of *Crellastrina* the most^67^.

Minimum age justification: Fossils were sampled from sediments that were collected from the outcrop that is part of the mid-Eocene to mid-Oligocene, 200 m thick succession that is overlayed by Pliocene and Pleistocene sediments^31^. More detailed fossilized spicules fall within the Priabonian age 33.9–37.71 Ma. They derived from two middle units of the Blanche Point Formation: Gull Rock member (Mb.) and Perkana Member.

**Node 103 Astrophorina** | 199 Ma

Fossil Taxon Specimen: Tetractinellida, Astrophorina (isolated dichotriaenes) Gabbs Valley Range of west central Nevada, USA.

Phylogenetic justification: arrays of long, complete, and complex in situ dichotriaenes of astrophorin demosponge affinity^32^ (see Ritterbush et al. 2015, Fig 6).

Minimum age justification: Fossils were found in the stratigraphic Unit of the Ferguson Hill Member of the Sunrise Formation within the Chert-dominated interval (24 to 55 m) which is of Hettangian and lower Sinemurian stages (199.3 Ma)^70^.

**Node 107: Crown Geodiidae** | 166.1-163.5 Ma

Fossil Taxon Specimen: *Geoditesia jordaniensis*, Holotype ESH 2009 I 34, a Callovian (Middle Jurassic) fossil from north-western Jordan.

Phylogenetic justification: well-preserved specimens of an articulated sponge belonging to Geodia due to the consisting of a homogeneous mass of intermingled spicules, mainly oxeas and triaenes; sterrasters are also present; inter structure is however, chaotic, but nevertheless, seem to represent recent morphological features of the genus *Geodia*^33^. We note that Cardenas (2020)^8^ has questioned the affiliation of this fossils, and we performed a sensitivity test where the calibrations where relaxed to consider this view (see Supplementary Methods).

Minimum age justification: Fossil is found in the Tal el Dhahab, Callovian marls of Mughanniyya Formation, Jordan which is of Callovian age 166.1–163.5 Ma (Middle Jurassic).

**Node 112 Spongillida** | 298 Ma

Fossil Taxon Specimen: Spongillida indet. PWL2004/5035a-LS, findspot Lemberg/Saar-Nahe Basin, layer 4, SW Germany.

Phylogenetic justification: morphological characters such as ordinary smooth monaxone spicules point to Spongillida. In addition, the fossils were from freshwater lake deposits placed far away from the sea, thus excluding a marine sponge assignment. However, gemmules and strongyles are absent, thus no fine scale classification can be made.

Minimum age justification: Permo-Carboniferous of Europe 298.9 Ma from the Saar-Nahe Basin (Stefanian C and Autunian) in south-west Germany. All fossils originated from freshwater lake deposits^34^.

**Node 128 Hexactinellida** | 445 Ma

Fossil Taxon Specimen: *Matteolaspongia hemiglobosa* (Holotype: NIGP168221). Found in the Anji Biota of the Wenchang Formation, middle to late *M. persculptus* Biozone, Hirnantian. Zhejiang Province, South China.

Phylogenetic justification: Based on morphological characterization (hypodermal pentactine prostalia, partly diactin-based skeleton) is fully consistent with a stem-Rossellidae interpretation, thus falls within hexactinellid sponges.

Minimum age justification: The fossil occurs in the latest Ordovician, the Hirnantian Age (445.2–443.8), more precisely dated to 444 Ma^35^.

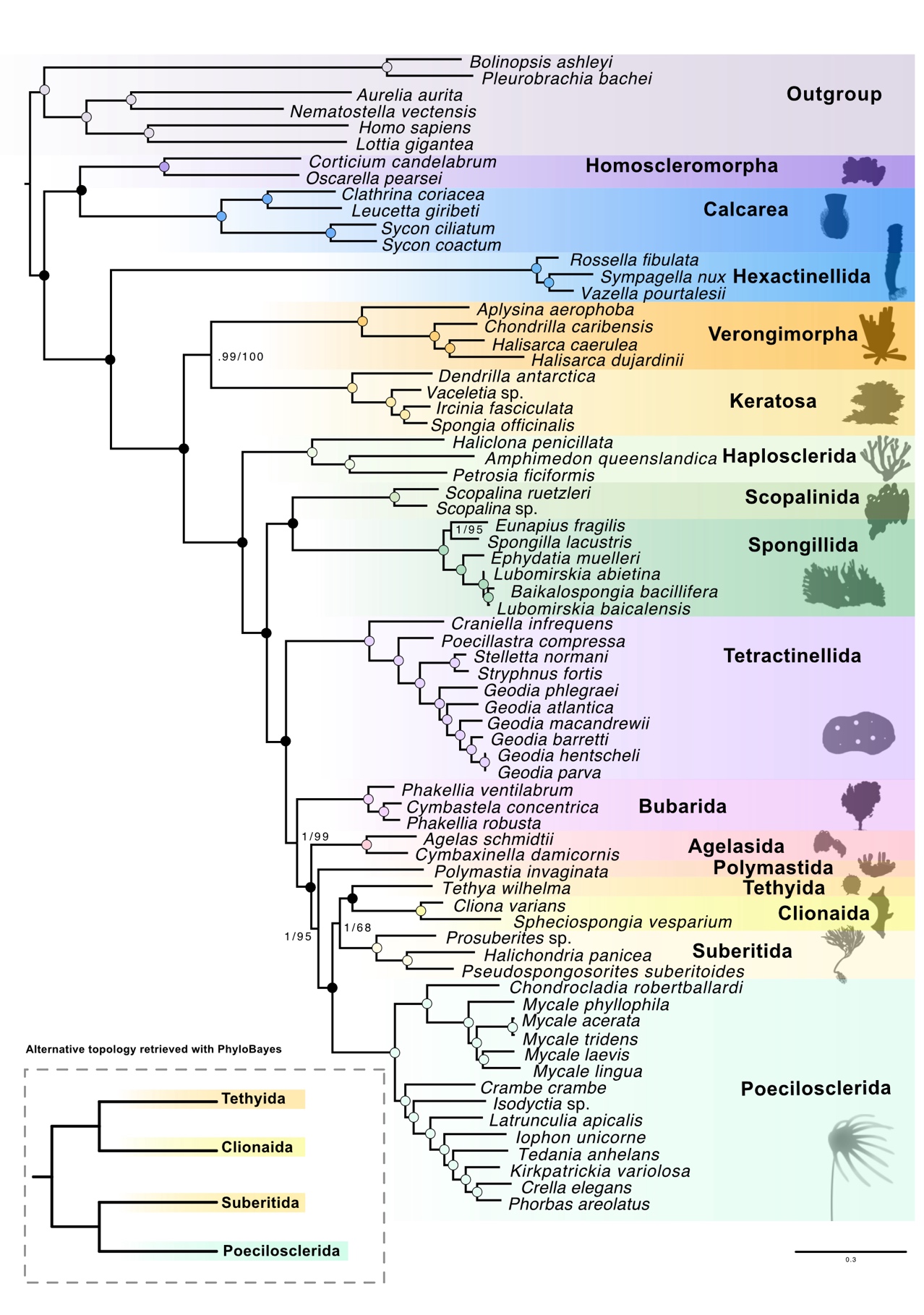

Fig. S1. Maximum likelihood topology of the phylum Porifera. Numbers on the nodes indicate posterior probability (BI) and bootstrap support (ML) respectively, fully supported nodes from both analyses are represented by full circles. In the grey box on the left, it is reported the alternative topology for Tethyida, Clionaida, Suberitida and Poecilosclerida from the Bayesian inference

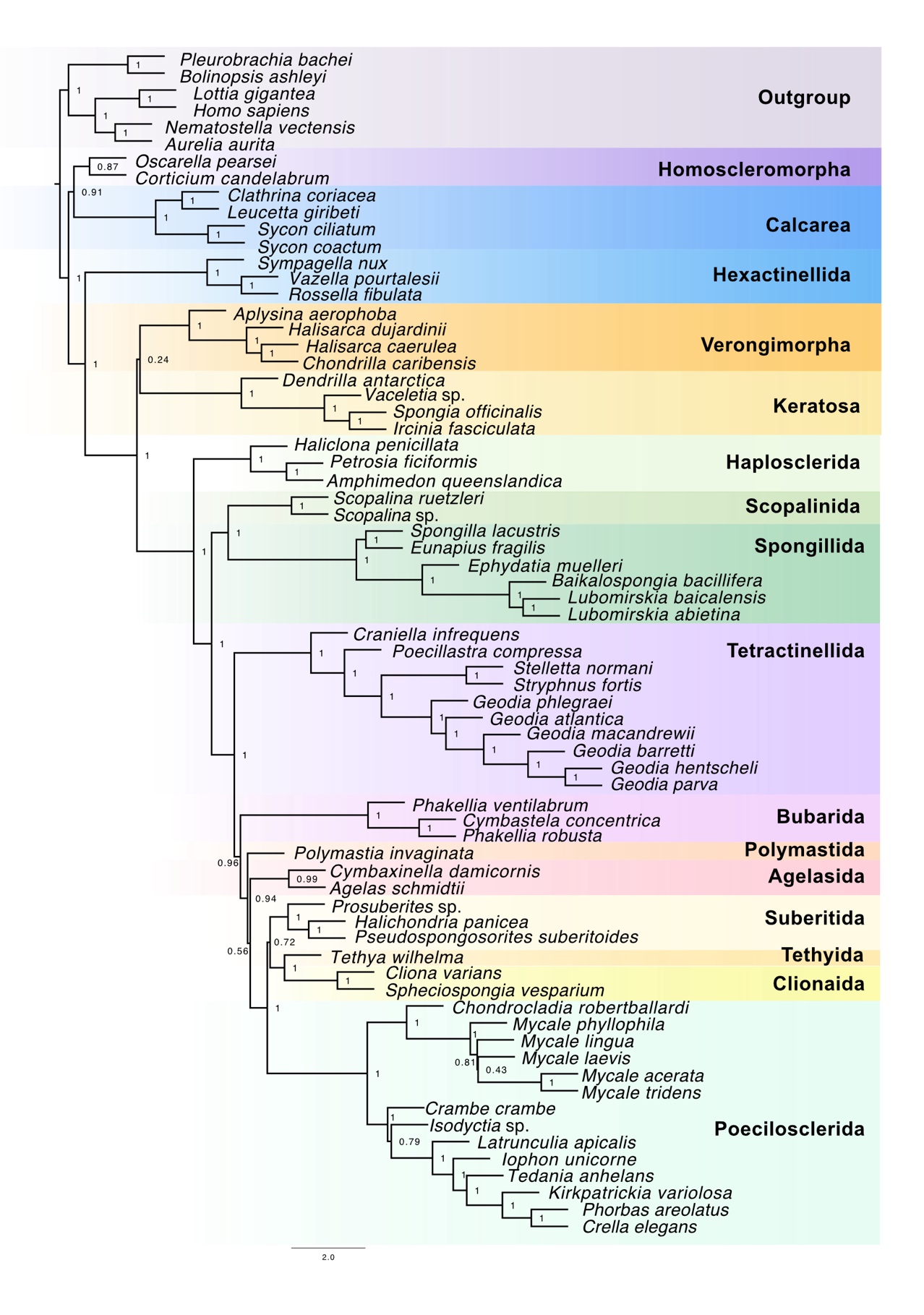

Fig. S2. Phylogenetic topology of Porifera inferred with ASTRAL. Number on the nodes indicates the coalescent branch support values.

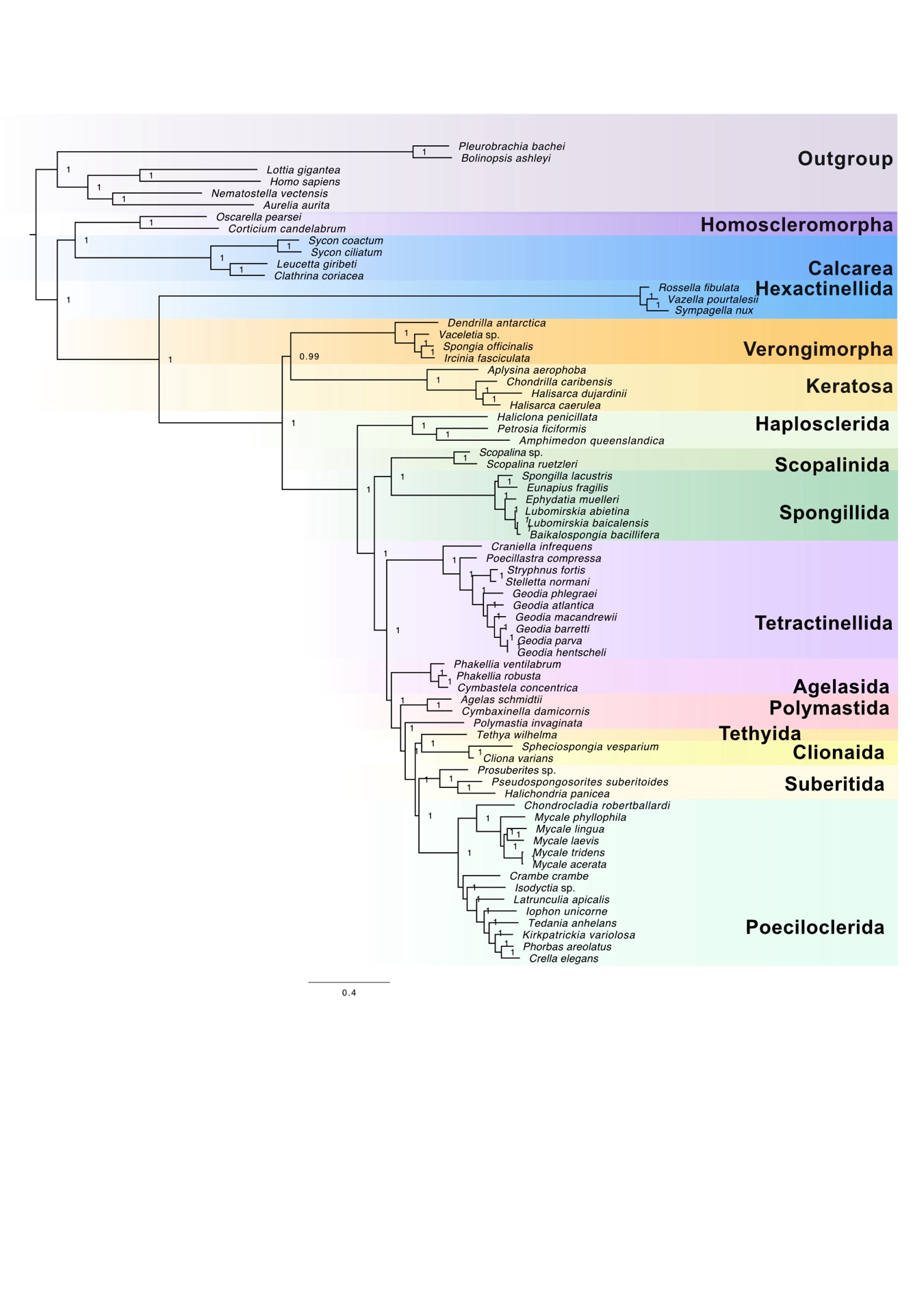

Fig. S3. Phylogenomic tree obtained with Phylobayes. Number on the nodes indicate the posterior probability values. Convergence was check with bpcomp (maxdiff = 0.0018315) and tracecomp (minimal ESS 61 and maximum real_diff=224488).

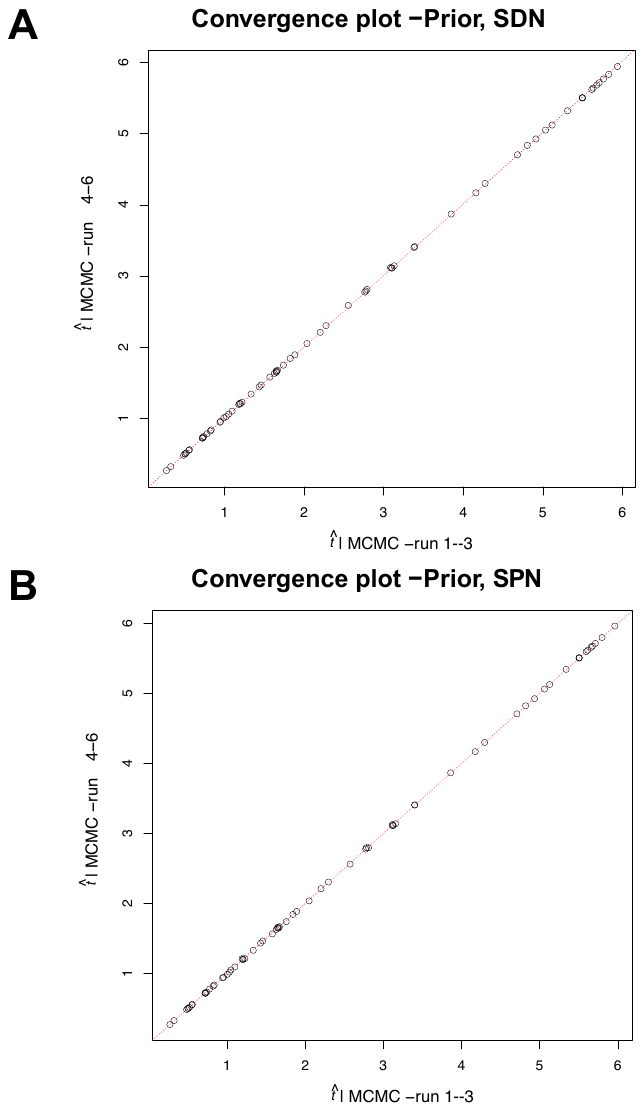

**Figure S4 Prior convergence** Convergence plot for the prior estimation of the two calibration strategies SPN and SDN

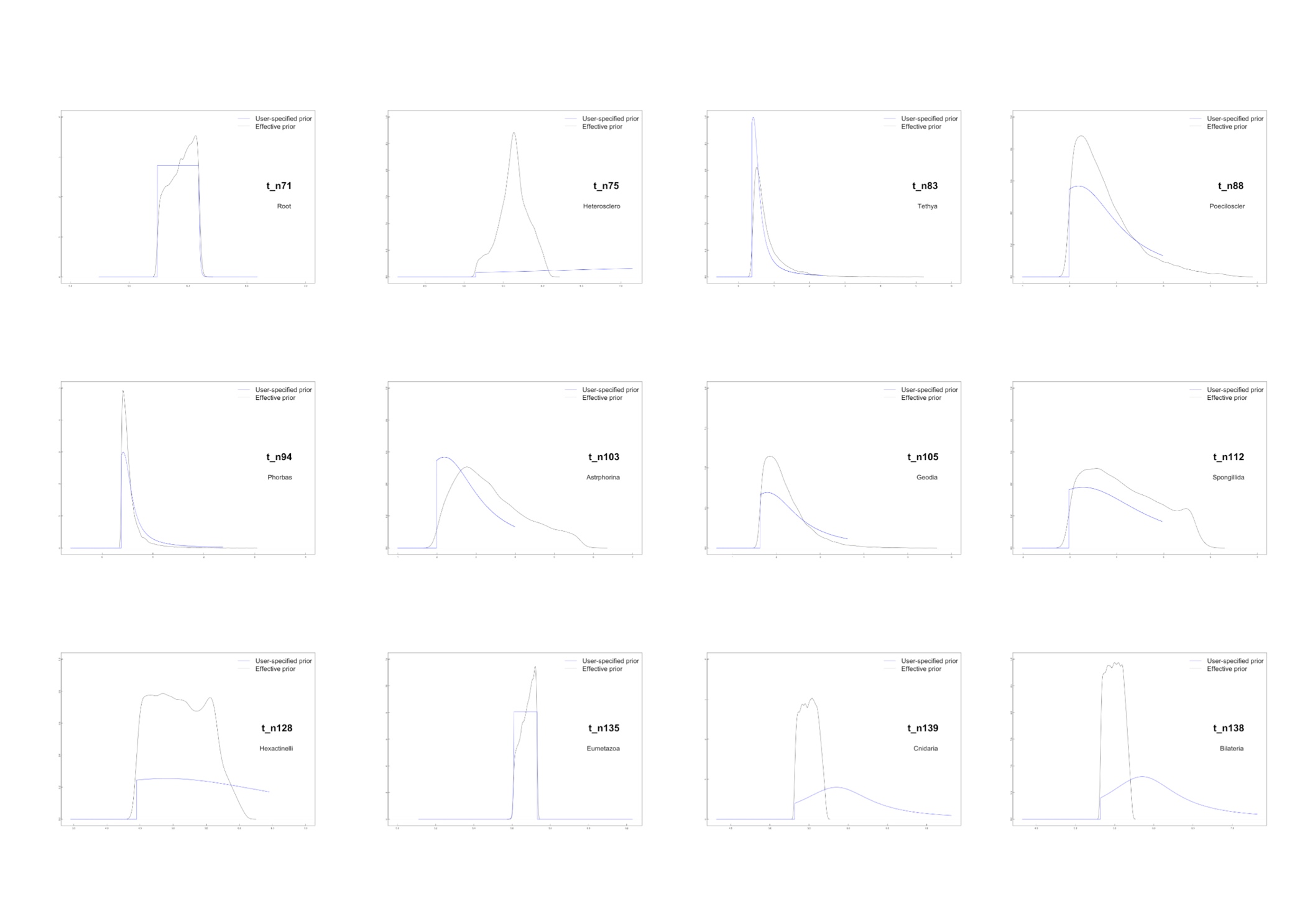

**Figure S5 Prior selection** Prior distribution plots of the calibration used for the divergence time estimates, comparing the user-specified prior (blue line) and the effective prior (black line) selected by MCMCTree.

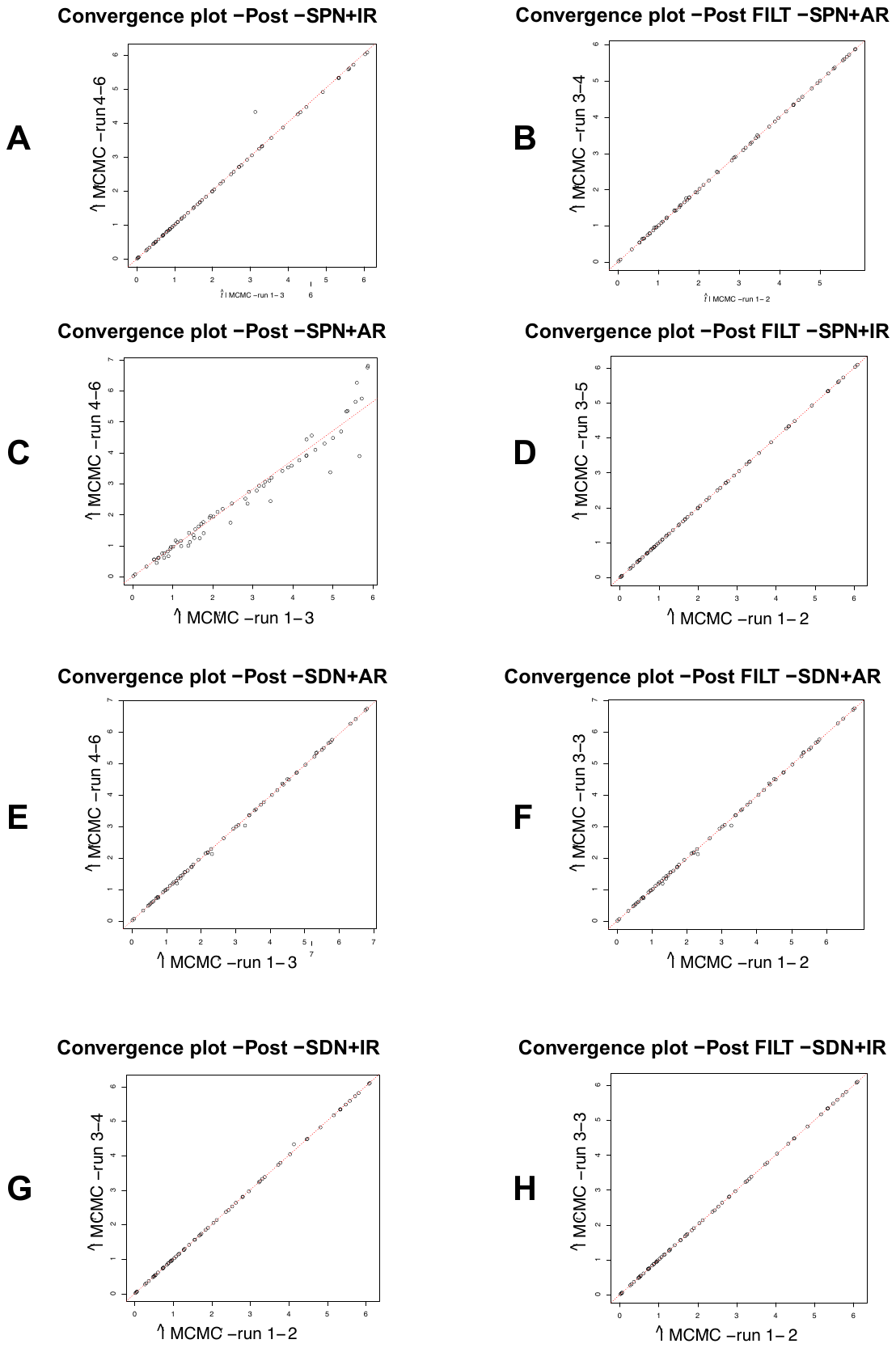

**Figure S6 Posterior convergence check** Convergence plots of the posterior time estimation for the two calibration strategies pre and post filtering for the chains flagged as problematic by the inhouse script MCMC diagnostics (see Methods).

**Figure S7** Porifera time-tree with posterior estimates using the SDN calibration strategy and the AR model in MCMCTree. Blue bars on nodes represent the 95% HPD. Time scale on x-axis in hundreds of millions of years. Abbreviation on the scale bar stands for Ne, Neoproterozoic; C, Cambrian; O, Ordovician; S, Silurian; D, Devonian; C, Carboniferous; P, Permian; Tr, Triassic; J, Jurassic; K, Cretaceous; Pe, Paleogene.

**Figure S8 SPN AR** Porifera time-tree with posterior estimates using the SPN calibration strategy and the AR model in MCMCTree. Blue bars on nodes represent the 95% HPD. Time scale on x-axis in hundreds of millions of years. Abbreviation on the scale bar stands for Ne, Neoproterozoic; C, Cambrian; O, Ordovician; S, Silurian; D, Devonian; C, Carboniferous; P, Permian; Tr, Triassic; J, Jurassic; K, Cretaceous; Pe, Paleogene.

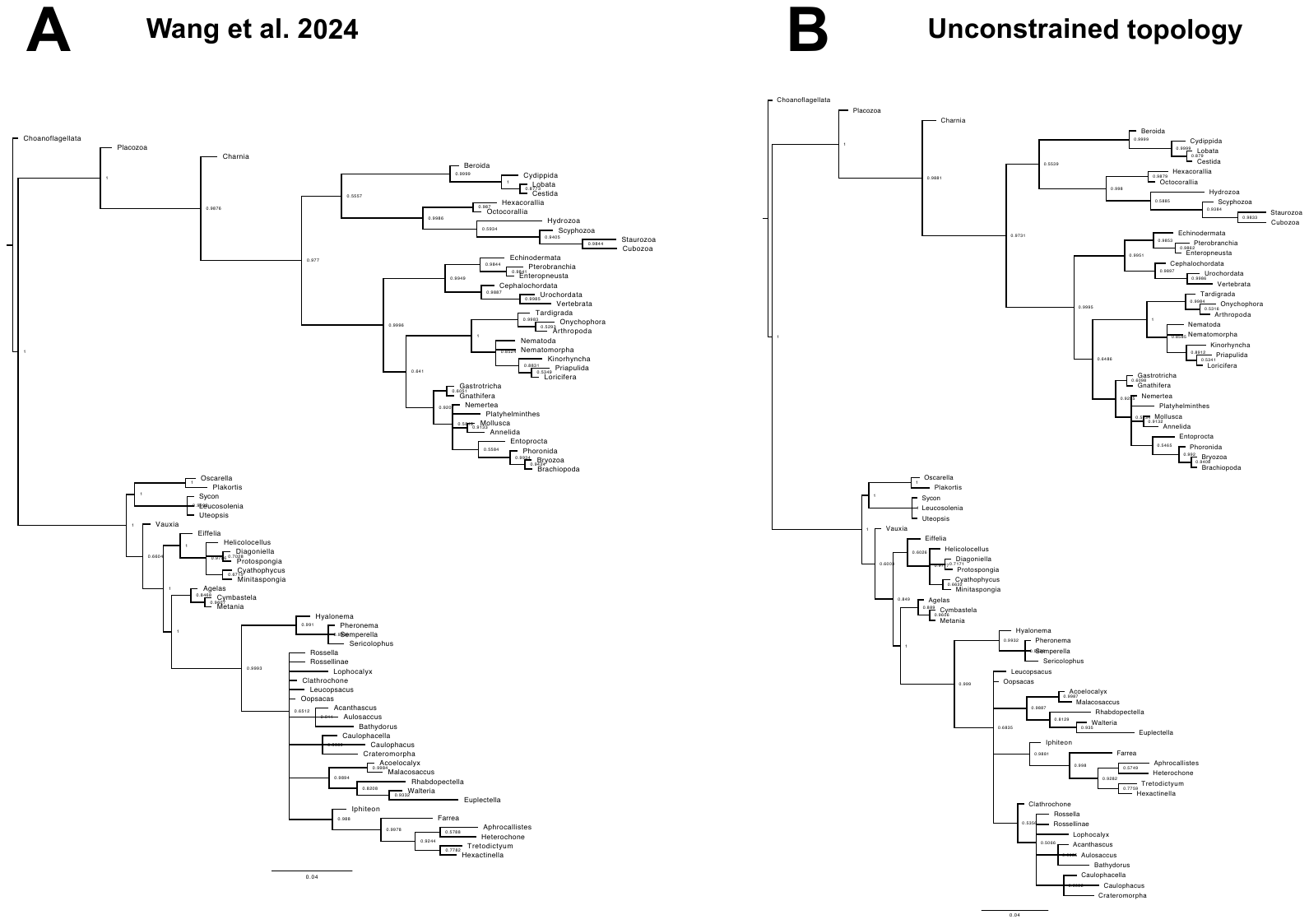

**Figure S10 (**A) Results of MrBayes analysis where extant taxa are constrained to follow results of our phylogenomic analyses and fossil taxa are constrained to belong to the same clade where they were found in Wang et al. original study. (B) Results of MrBayes analyses where the extant taxa were constrained to follow the results of our phylogenomic analyses and the fossil taxa were free to place in their optimal (unconstrained) position in the tree topology.

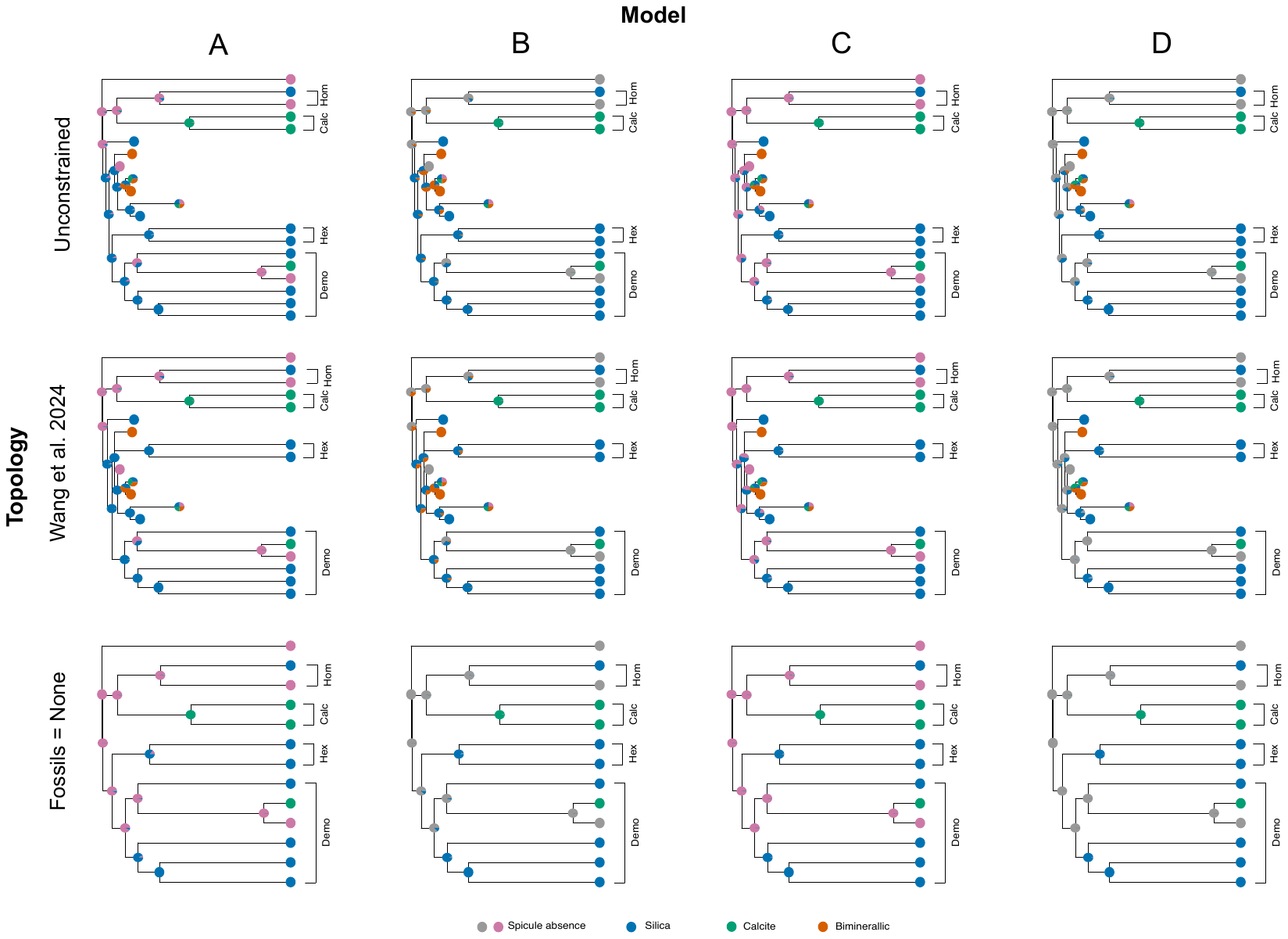

**Figure S11** Simplified summary results of the ancestral state estimations. Horizontally the trees are grouped by the topologies, and vertically by models. The legend depict each color assigned to mineralogy. Pink/Grey absence of spicules, Blue for siliceous spicules, Green for calcareous spicules, and Orange for biminerallic spicules. Here we show only mineralogies, but not biochemical differences in the developmental of spicules of a specific type. To see full results please refer to <https://github.com/MEleonoraRossi/ASE_spicules/>

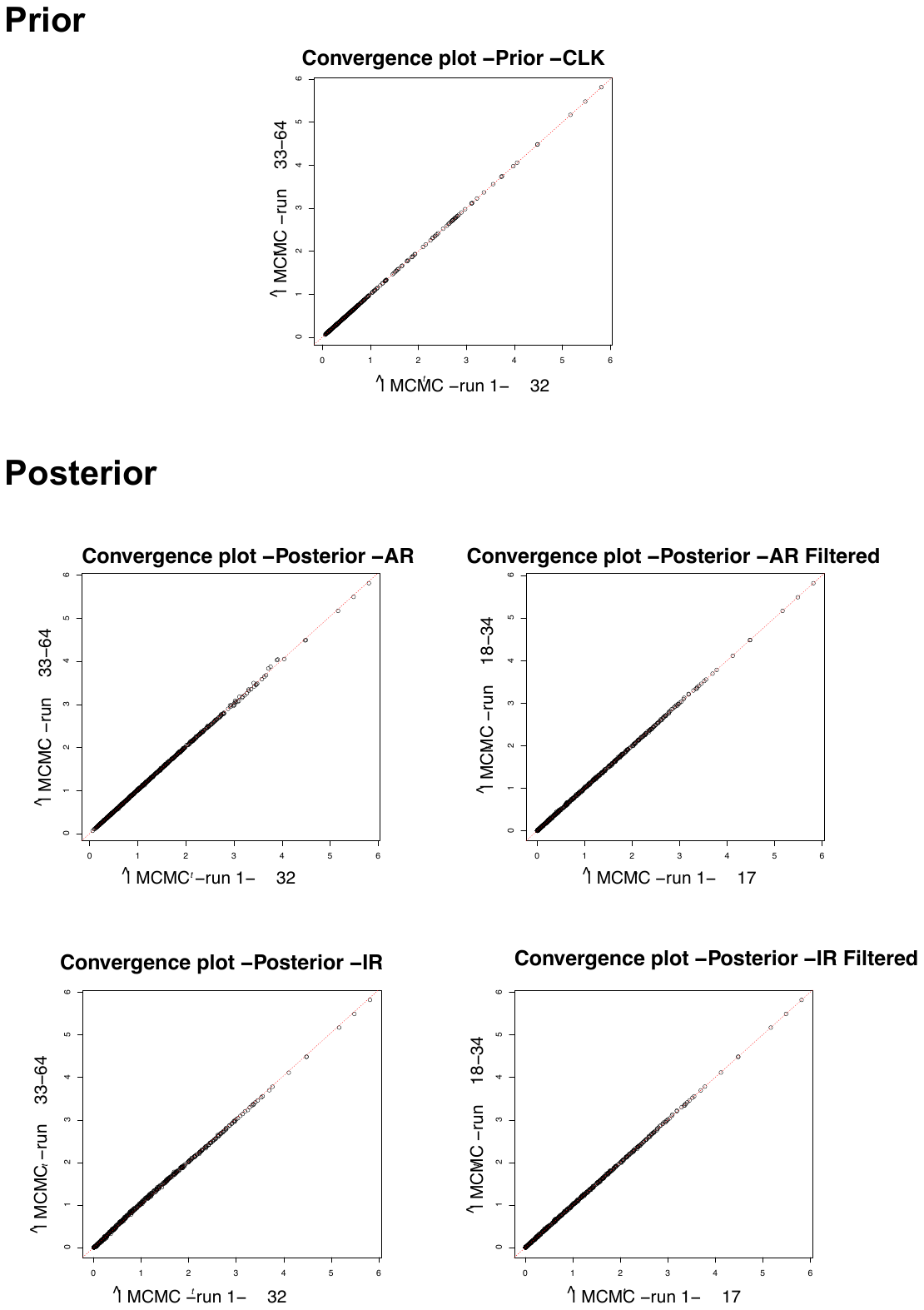

**Figure S12** Convergence plots of the posterior time estimation for the COI topology used for diversification rates analyses, with pre and post filtering for the chains flagged as problematic by our in-house script MCMC diagnostics.

Table S1.

**Table S1** MCMC diagnostics for time densities obtained with MCMCtree when fixing the SDN tree topology, both when the target distribution was set to be the prior (no data, label “CLK”) or the posterior (data were used, labels for relaxed-clock models are “AR” for autocorrelated rates and “IR” for independent rates sampled from a log-normal distribution).

| SDN Strategy |  |  |  |
| --- | --- | --- | --- |
|  | **Prior-CLK** | **Autocorrelated-AR** | **Independent-IR** |
| tail-ESS times (median) | 30,865 | 1,063 | 3,374 |
| tail-ESS times (min) | 3,359 | 273 | 686 |
| tail-ESS times (max) | 53,466 | 23,413 | 28,471 |
| bulk-ESS times (median) | 33,260 | 667 | 1873 |
| bulk-ESS times (min) | 6,695 | 133 | 348 |
| bulk-ESS times (max) | 45,726 | 18,287 | 29,085 |
| Rhat min | 0.9999812 | 0.9999731 | 0.9999337 |
| Rhat max | 1.001506 | 1.038014 | 1.006146 |
| Med. number of samples per chain | 20,001 | 20,001 | 20,001 |
| Min. number of samples per chain | 20,001 | 20,001 | 20,001 |
| Max. number of samples per chain | 20,001 | 20,001 | 20,001 |
| Number of chains that were run | 6 | 6 | 6 |
| Number of chains kept after filters | 6 | 3 | 3 |
| Number of samples used to calculate tail-ESS and bulk-ESS | 120,006 | 60,003 | 60,003 |
| Total number of samples kept *post-hoc* analyses | 120,006 | 60,003 | 60,003 |

**Table S2** MCMC diagnostics for time densities obtained with MCMCtree when fixing the SPN tree topology, both when the target distribution was set to be the prior (no data, label “CLK”) or the posterior (data were used, labels for relaxed-clock models are “AR” for autocorrelated rates and “IR” for independent rates sampled from a log-normal distribution).

| SPN strategy |  |  |  |
| --- | --- | --- | --- |
|  | **Prior-CLK** | **Autocorrelated- AR** | **Independent-IR** |
| tail-ESS times (median) | 30,662 | 1,348 | 6,014 |
| tail-ESS times (min) | 3,661 | 411 | 1555 |
| tail-ESS times (max) | 59,697 | 33,096 | 49,641 |
| bulk-ESS times (median) | 33,336 | 741 | 3197 |
| bulk-ESS times (min) | 6,451 | 214 | 768 |
| bulk-ESS times (max) | 58,044 | 29,179 | 46,930 |
| Rhat min | 0.999989 | 1.000045 | 0.9999561 |
| Rhat max | 1.000565 | 1.025086 | 1.005087 |
| Med. number of samples per chain | 20,001 | 20,001 | 20,001 |
| Min. number of samples per chain | 20,001 | 20,001 | 20,001 |
| Max. number of samples per chain | 20,001 | 20,001 | 20,001 |
| Number of chains that were run | 6 | 6 | 6 |
| Number of chains kept after filters | 6 | 4 | 5 |
| Number of samples used to calculate tail-ESS and bulk-ESS | 120,006 | 80,004 | 100,005 |
| Total number of samples kept *post-hoc* analyses | 120,006 | 80,004 | 100,005 |

**Table S3** Mean posterior ivergence times estimated wit MCMCtree when the calibration strategy was SPN under  both relaxed-clock models (“AR” for autocorrelated rates and “IR” for independent rates sampled from a log-normal distribution).

| **SPN-IR** | | | | | **SPN-AR** | | | | |
| --- | --- | --- | --- | --- | --- | --- | --- | --- | --- |
|  | **Mean div. time** | **2.5% CI** | **97.5% CI** | **Priors** |  | **Mean div. time** | **2.5% CI** | **97.5% CI** | **Priors** |
| t_n71 ROOT | 608.582 | 603.28 | 613.15 | B(5.7400,6.0900,1e-300,0.0250) | T_n71 ROOT | 587.641 | 581.39 | 595.25 | B(5.7400,6.0900,1e-300,0.0250) |
| t_n72 PORIFERA | 602.821 | 590.34 | 610.68 | L(5.1500,0.5000,0.5000,1e-300) | t_n72 PORIFERA | 586.586 | 578.79 | 594.59 | L(5.1500,0.5000,0.5000,1e-300) |
| t_n73 | 561.092 | 542.68 | 575.87 | 0 | t_n73 | 559.951 | 551.55 | 567.69 | 0 |
| t_n74 | 491.825 | 462.67 | 519.67 | 0 | t_n74 | 520.579 | 503.28 | 535.02 | 0 |
| t_n75 | 426.204 | 396.53 | 454.95 | 0 | t_n75 | 479.364 | 455.96 | 500.69 | 0 |
| t_n76 | 387.255 | 359.79 | 414.91 | 0 | t_n76 | 456.313 | 431.08 | 480.01 | 0 |
| t_n77 | 356.451 | 330.46 | 383.55 | 0 | t_n77 | 434.377 | 407.97 | 459.28 | 0 |
| t_n78 | 332.509 | 307.43 | 359.25 | 0 | t_n78 | 416.176 | 388.49 | 442.35 | 0 |
| t_n79 | 305.303 | 282.35 | 330.66 | 0 | t_n79 | 388.33 | 357.39 | 417.34 | 0 |
| t_n80 | 120.702 | 57.97 | 202.37 | 0 | t_n80 | 202.386 | 99.8 | 308.88 | 0 |
| t_n81 | 292.378 | 270.1 | 317.26 | 0 | t_n81 | 374.188 | 342.07 | 404.57 | 0 |
| t_n82 | 271.005 | 250.84 | 294.45 | 0 | t_n82 | 347.02 | 313.96 | 379.31 | 0 |
| t_n83 THETHYA | 228.79 | 186.07 | 263.73 | L(0.3780,0.1000,0.5000,1e-300) | t_n83 THETHYA | 315.936 | 281.05 | 351.02 | L(0.3780,0.1000,0.5000,1e-300) |
| t_n84 | 45.963 | 23 | 82.93 | 0 | t_n84 | 108.33 | 53.71 | 174.79 | 0 |
| t_n85 | 256.913 | 237.85 | 279.57 | 0 | t_n85 | 330.854 | 297.89 | 364.01 | 0 |
| t_n86 | 173.353 | 130.76 | 213.96 | 0 | t_n86 | 247.972 | 186.65 | 299.89 | 0 |
| t_n87 | 108.812 | 75.22 | 146.6 | 0 | t_n87 | 165.207 | 112.14 | 218.35 | 0 |
| t_n88 POECILOSCLERIDA | 205.294 | 199.21 | 219.24 | L(1.9900,0.1000,0.5000,1e-300) | t_n88  POECILOSCLERIDA | 212.59 | 199.41 | 240.17 | L(1.9900,0.1000,0.5000,1e-300) |
| t_n89 | 183.281 | 166.16 | 199.54 | 0 | t_n89 | 192.825 | 176.82 | 220.16 | 0 |
| t_n90 | 167.081 | 150.1 | 184.3 | 0 | t_n90 | 177.225 | 159.44 | 204.31 | 0 |
| t_n91 | 135.932 | 117.78 | 154.46 | 0 | t_n91 | 141.589 | 119.98 | 168.11 | 0 |
| t_n92 | 109.163 | 91.81 | 127.4 | 0 | t_n92 | 111.74 | 90.5 | 136.84 | 0 |
| t_n93 | 93.894 | 77.43 | 111.69 | 0 | t_n93 | 96.071 | 76.51 | 119.41 | 0 |
| t_n94 PHORBAS | 70.024 | 54.79 | 87.4 | L(0.3780,0.1000,0.5000,1e-300) | t_n94 PHORBAS | 74.291 | 57.61 | 94.9 | L(0.3780,0.1000,0.5000,1e-300) |
| t_n95 | 49.089 | 35.03 | 65.46 | 0 | t_n95 | 54.232 | 40.57 | 71.68 | 0 |
| t_n96 | 149.998 | 126.43 | 171.73 | 0 | t_n96 | 157.111 | 134.25 | 185.67 | 0 |
| t_n97 | 88.803 | 69.76 | 109.63 | 0 | t_n97 | 94.913 | 72.38 | 120.65 | 0 |
| t_n98 | 71.422 | 53.88 | 91.33 | 0 | t_n98 | 79.37 | 59.08 | 102.81 | 0 |
| t_n99 | 58.179 | 41 | 78.07 | 0 | t_n99 | 64.237 | 46.69 | 85.04 | 0 |
| t_n100 | 5.358 | 3.31 | 8.22 | 0 | t_n100 | 6.994 | 4 | 11.56 | 0 |
| t_n101 | 68.219 | 48.26 | 94.21 | 0 | t_n101 | 143.841 | 65.09 | 230.21 | 0 |
| t_n102 | 34.13 | 21.91 | 50.78 | 0 | t_n102 | 79.325 | 32.34 | 141.67 | 0 |
| t_n103 ASTROPHORINA | 249.583 | 224.9 | 277.02 | L(1.9900,0.1000,0.5000,1e-300) | t_n103  ASTROPHORINA | 290.797 | 257.3 | 330.42 | L(1.9900,0.1000,0.5000,1e-300) |
| t_n104 | 199.961 | 185.43 | 219.12 | 0 | t_n104 | 225.269 | 203.77 | 256.23 | 0 |
| t_n105 GEODIA | 166.7 | 163.1 | 176.04 | L(1.6300,0.1000,0.5000,1e-300) | t_n105 GEODIA | 171.826 | 163.22 | 193.93 | L(1.6300,0.1000,0.5000,1e-300) |
| t_n106 | 28.366 | 17.61 | 43.29 | 0 | t_n106 | 54.084 | 32.7 | 80.15 | 0 |
| t_n107 | 103.253 | 81.47 | 126.24 | 0 | t_n107 | 120.904 | 101.35 | 142.95 | 0 |
| t_n108 | 84.37 | 64.9 | 106.33 | 0 | t_n108 | 101.644 | 82.52 | 122.86 | 0 |
| t_n109 | 51.263 | 37.5 | 67.97 | 0 | t_n109 | 65.623 | 49.6 | 84 | 0 |
| t_n110 | 25.478 | 16.86 | 36.71 | 0 | t_n110 | 35.189 | 24.76 | 48.29 | 0 |
| t_n111 | 1.788 | 1.01 | 2.93 | 0 | t_n111 | 2.232 | 1.26 | 3.74 | 0 |
| t_n112 SPONGILLIDA | 330.911 | 301.03 | 366.72 | L(2.9800,0.1000,0.5000,1e-300) | t_n112  SPONGILLIDA | 433.722 | 407.5 | 459.41 | L(2.9800,0.1000,0.5000,1e-300) |
| t_n113 | 80.812 | 49.96 | 123.92 | 0 | t_n113 | 194.3 | 106.81 | 306.48 | 0 |
| t_n114 | 87.71 | 66.94 | 113.02 | 0 | t_n114 | 327.482 | 294.99 | 359.8 | 0 |
| t_n115 | 70.025 | 49.56 | 95.19 | 0 | t_n115 | 309.857 | 275.13 | 344.41 | 0 |
| t_n116 | 44.267 | 29.96 | 62.55 | 0 | t_n116 | 281.559 | 247.13 | 316.9 | 0 |
| t_n117 | 5.292 | 3.73 | 7.29 | 0 | t_n117 | 152.267 | 124.47 | 183.33 | 0 |
| t_n118 | 3.415 | 2.25 | 4.93 | 0 | t_n118 | 87.932 | 64.23 | 116.5 | 0 |
| t_n119 | 276.571 | 213.8 | 336.29 | 0 | t_n119 | 397.502 | 338.15 | 444.03 | 0 |
| t_n120 | 198.845 | 142.11 | 263.92 | 0 | t_n120 | 341.744 | 265.18 | 404.88 | 0 |
| t_n121 | 432.272 | 390.96 | 470.4 | 0 | t_n121 | 500.244 | 477.51 | 519.44 | 0 |
| t_n122 | 161.414 | 122.02 | 209.26 | 0 | t_n122 | 172.572 | 69.01 | 313.43 | 0 |
| t_n123 | 78.792 | 57.86 | 104.96 | 0 | t_n123 | 92.724 | 33.67 | 187.79 | 0 |
| t_n124 | 50.749 | 33.8 | 72.88 | 0 | t_n124 | 62.607 | 21.75 | 133.52 | 0 |
| t_n125 | 221.766 | 169.77 | 281.75 | 0 | t_n125 | 287.991 | 177.67 | 373.04 | 0 |
| t_n126 | 118.131 | 87.8 | 154.84 | 0 | t_n126 | 155.17 | 85.78 | 227.96 | 0 |
| t_n127 | 88.413 | 61.29 | 122.31 | 0 | t_n127 | 122.447 | 65.41 | 188.14 | 0 |
| t_n128 HEXACTINELLIDA | 447.732 | 445.07 | 454.99 | L(4.4500,0.1000,0.5000,1e-300) | T_n128 HEXACTINELLIDA | 447.308 | 445.06 | 453.36 | L(4.4500,0.1000,0.5000,1e-300) |
| t_n129 | 433.117 | 415.1 | 444.85 | 0 | t_n129 | 434.484 | 422.5 | 443.89 | 0 |
| t_n130 | 532.548 | 480.93 | 562.42 | 0 | t_n130 | 566.379 | 556.69 | 574.07 | 0 |
| t_n131 | 324.35 | 224.87 | 421.58 | 0 | t_n131 | 493.625 | 420.18 | 535.63 | 0 |
| t_n132 | 271.411 | 207.53 | 337.36 | 0 | t_n132 | 347.621 | 157.52 | 460.38 | 0 |
| t_n133 | 97.804 | 63.93 | 141.51 | 0 | t_n133 | 141.306 | 49.76 | 264.43 | 0 |
| t_n134 | 152.246 | 102.27 | 208.33 | 0 | t_n134 | 247.368 | 96.14 | 381.18 | 0 |
| t_n135 EUMETAZOA | 572.198 | 569.91 | 573.08 | B(5.6100,5.7310,0.0250,1e-300) | T_n135 EUMETAZOA | 572.138 | 569.63 | 573.08 | B(5.6100,5.7310,0.0250,1e-300) |
| t_n136 | 125.961 | 72.69 | 195.46 | 0 | t_n136 | 178.163 | 65.34 | 352.78 | 0 |
| t_n137 | 558.524 | 554.03 | 562.73 | 0 | t_n137 | 556.568 | 553.43 | 559.54 | 0 |
| t_n138 BILATERIA | 533.43 | 532.04 | 537.03 | L(5.3200,0.1000,0.1000,1e-300) | T_n138 BILATERIA | 533.696 | 532.05 | 537.65 | L(5.3200,0.1000,0.1000,1e-300) |
| t_n139 CNIDARIA | 533.492 | 531.85 | 537.66 | L(5.3180,0.1000,0.1000,1e-300) | T_n139 CNIDARIA | 536.527 | 532.1 | 542.54 | L(5.3180,0.1000,0.1000,1e-300) |

Table S4  Divergence time estimates obtained with the calibration strategy SDN, for both molecular clocks IR and AR

| **SDN-ILN** | | |  | | | **SDN-AR** | | |  | | |
| --- | --- | --- | --- | --- | --- | --- | --- | --- | --- | --- | --- |
|  | **Mean div. time** | **2.5% CI** | | **97.5% CI** | **Priors** |  | **Mean div. time** | **2.5% CI** | | **97.5% CI** | **Priors** |
| T_n71 ROOT | 610.315 | 605.92 | | 616.91 | B(5.7400,6.0900,1e-300,0.0250) | T_n71 ROOT | 591.484 | 583.49 | | 604.27 | B(5.7400,6.0900,1e-300,0.0250) |
| t_n72 | 608.138 | 601.3 | | 615.32 | 0 | t_n72 | 590.996 | 582.68 | | 603.79 | 0 |
| t_n73 | 581.288 | 570.3 | | 591.32 | 0 | t_n73 | 568.471 | 561.72 | | 577.48 | 0 |
| t_n74 | 547.883 | 536.53 | | 559.17 | 0 | t_n74 | 543.705 | 538.18 | | 549.57 | 0 |
| T_n75  HETEROSCLEROMORPHA | 516.905 | 515.05 | | 521.97 | L(5.1500,0.5000,0.5000,1e-300) | T_n75  HETEROSCLEROMORPHA | 516.689 | 515.04 | | 520.81 | L(5.1500,0.5000,0.5000,1e-300) |
| t_n76 | 447.309 | 412.61 | | 478.57 | 0 | t_n76 | 496.554 | 489.5 | | 503.17 | 0 |
| t_n77 | 403.951 | 368.75 | | 438.04 | 0 | t_n77 | 474.376 | 460.89 | | 485.45 | 0 |
| t_n78 | 373.428 | 338.39 | | 408.31 | 0 | t_n78 | 455.511 | 436.28 | | 470.99 | 0 |
| t_n79 | 338.08 | 305.31 | | 372.19 | 0 | t_n79 | 425.887 | 397.75 | | 448.66 | 0 |
| t_n80 | 126.527 | 59.98 | | 215.67 | 0 | t_n80 | 223.256 | 104.76 | | 346.79 | 0 |
| t_n81 | 322.887 | 290.87 | | 356.77 | 0 | t_n81 | 410.679 | 379.4 | | 436.63 | 0 |
| t_n82 | 296.733 | 267.19 | | 330.03 | 0 | t_n82 | 380.923 | 345.13 | | 412.03 | 0 |
| T_n83 THETHYA | 252.87 | 199.04 | | 295.9 | L(0.3780,0.1000,0.5000,1e-300) | T_n83  THETHYA | 347.102 | 306.93 | | 383.57 | L(0.3780,0.1000,0.5000,1e-300) |
| t_n84 | 49.452 | 24.03 | | 91.25 | 0 | t_n84 | 121.145 | 58.61 | | 197.27 | 0 |
| t_n85 | 280.107 | 251.82 | | 312.48 | 0 | t_n85 | 363.014 | 325.49 | | 396.69 | 0 |
| t_n86 | 184.903 | 136.41 | | 236.04 | 0 | t_n86 | 271.878 | 194.47 | | 330.76 | 0 |
| t_n87 | 115.62 | 78.06 | | 158.71 | 0 | t_n87 | 180.543 | 117.84 | | 243.11 | 0 |
| T_n88  POECILOSCLERIDA | 213.195 | 199.74 | | 236.53 | L(1.9900,0.1000,0.5000,1e-300) | T_n88  POECILOSCLERIDA | 218.2 | 199.63 | | 254.93 | L(1.9900,0.1000,0.5000,1e-300) |
| t_n89 | 190.382 | 169.92 | | 214.39 | 0 | t_n89 | 197.508 | 176.93 | | 233.34 | 0 |
| t_n90 | 173.422 | 153 | | 197.6 | 0 | t_n90 | 181.442 | 159.5 | | 216.33 | 0 |
| t_n91 | 141.298 | 120.69 | | 163.96 | 0 | t_n91 | 144.91 | 119.84 | | 177.89 | 0 |
| t_n92 | 113.475 | 94.44 | | 134.56 | 0 | t_n92 | 114.52 | 90.14 | | 144.97 | 0 |
| t_n93 | 97.572 | 79.51 | | 118.07 | 0 | t_n93 | 98.592 | 76.25 | | 126.6 | 0 |
| T_n94  PHORBAS | 72.542 | 56.28 | | 92.04 | L(0.3780,0.1000,0.5000,1e-300) | T_n94  PHORBAS | 76.423 | 57.51 | | 100.36 | L(0.3780,0.1000,0.5000,1e-300) |
| t_n95 | 50.735 | 35.93 | | 68.78 | 0 | t_n95 | 55.927 | 40.67 | | 75.77 | 0 |
| t_n96 | 155.672 | 130 | | 182.52 | 0 | t_n96 | 160.508 | 133.89 | | 195.95 | 0 |
| t_n97 | 93.13 | 72.97 | | 115.65 | 0 | t_n97 | 96.756 | 71.98 | | 127.16 | 0 |
| t_n98 | 74.793 | 55.94 | | 96.3 | 0 | t_n98 | 80.945 | 58.82 | | 108.45 | 0 |
| t_n99 | 60.986 | 42.52 | | 82.7 | 0 | t_n99 | 65.431 | 46.41 | | 89.64 | 0 |
| t_n100 | 5.695 | 3.43 | | 8.9 | 0 | t_n100 | 7.208 | 4.02 | | 12.31 | 0 |
| t_n101 | 73.476 | 51.51 | | 103.05 | 0 | t_n101 | 155.794 | 65.96 | | 256.3 | 0 |
| t_n102 | 36.565 | 23.06 | | 55.3 | 0 | t_n102 | 86.468 | 32.82 | | 160.87 | 0 |
| T_n103 ASTROPHORINA | 263.023 | 231.07 | | 299.37 | L(1.9900,0.1000,0.5000,1e-300) | T_n103 ASTROPHORINA | 302.114 | 263.11 | | 349.6 | L(1.9900,0.1000,0.5000,1e-300) |
| t_n104 | 204.745 | 187.39 | | 228.83 | 0 | t_n104 | 229.253 | 205.57 | | 265.07 | 0 |
| T_n105 GEODIA | 167.654 | 163.13 | | 179.28 | L(1.6300,0.1000,0.5000,1e-300) | t_n105_GEODIA | 172.998 | 163.27 | | 197.64 | L(1.6300,0.1000,0.5000,1e-300) |
| t_n106 | 30.069 | 18.53 | | 46.3 | 0 | t_n106 | 54.279 | 32.78 | | 80.31 | 0 |
| t_n107 | 107.462 | 85.01 | | 130.04 | 0 | t_n107 | 121.203 | 101.4 | | 144.61 | 0 |
| t_n108 | 87.836 | 67.81 | | 110.28 | 0 | t_n108 | 101.817 | 82.55 | | 124.18 | 0 |
| t_n109 | 53.773 | 39.25 | | 71.69 | 0 | t_n109 | 65.656 | 49.55 | | 84.89 | 0 |
| t_n110 | 26.775 | 17.55 | | 38.97 | 0 | t_n110 | 35.239 | 24.75 | | 48.81 | 0 |
| t_n111 | 1.888 | 1.04 | | 3.12 | 0 | t_n111 | 2.236 | 1.25 | | 3.79 | 0 |
| T_n112  SPONGILLIDA | 378.697 | 325.85 | | 424.69 | L(2.9800,0.1000,0.5000,1e-300) | T_n112  SPONGILLIDA | 475.044 | 463 | | 485.34 | L(2.9800,0.1000,0.5000,1e-300) |
| t_n113 | 87.099 | 52.42 | | 135.13 | 0 | t_n113 | 218.685 | 114.69 | | 345.78 | 0 |
| t_n114 | 94.741 | 71.52 | | 123.21 | 0 | t_n114 | 363.38 | 333.66 | | 390.36 | 0 |
| t_n115 | 75.801 | 52.9 | | 103.46 | 0 | t_n115 | 345.117 | 311.74 | | 375.54 | 0 |
| t_n116 | 47.652 | 31.88 | | 68.43 | 0 | t_n116 | 314.379 | 279.92 | | 347.62 | 0 |
| t_n117 | 5.56 | 3.86 | | 7.72 | 0 | t_n117 | 170.195 | 139.74 | | 203.04 | 0 |
| t_n118 | 3.619 | 2.35 | | 5.3 | 0 | t_n118 | 100.024 | 72.96 | | 132.72 | 0 |
| t_n119 | 326.227 | 246.5 | | 403.84 | 0 | t_n119 | 450.517 | 409.36 | | 475.43 | 0 |
| t_n120 | 237.875 | 159.76 | | 321.1 | 0 | t_n120 | 401.213 | 338.32 | | 445.46 | 0 |
| t_n121 | 482.105 | 430.44 | | 521.85 | 0 | t_n121 | 530.528 | 523.55 | | 536.22 | 0 |
| t_n122 | 170.707 | 126.91 | | 227.91 | 0 | t_n122 | 162.39 | 65.48 | | 319.51 | 0 |
| t_n123 | 83.447 | 60.14 | | 112.7 | 0 | t_n123 | 86.319 | 32.09 | | 186.12 | 0 |
| t_n124 | 53.595 | 34.93 | | 78.44 | 0 | t_n124 | 58.064 | 20.7 | | 130.74 | 0 |
| t_n125 | 242.798 | 180.75 | | 310.25 | 0 | t_n125 | 319.44 | 181.23 | | 415.25 | 0 |
| t_n126 | 127.479 | 92.17 | | 168.66 | 0 | t_n126 | 173.153 | 87.17 | | 263.53 | 0 |
| t_n127 | 95.455 | 64.34 | | 133.71 | 0 | t_n127 | 137.371 | 66.35 | | 218.44 | 0 |
| T_n128  HEXACTINELLIDA | 448.388 | 445.09 | | 457.3 | L(4.4500,0.1000,0.5000,1e-300) | T_n128  HEXACTINELLIDA | 447.771 | 445.07 | | 454.92 | L(4.4500,0.1000,0.5000,1e-300) |
| t_n129 | 432.639 | 411.66 | | 446.13 | 0 | t_n129 | 433.521 | 419.64 | | 444.51 | 0 |
| t_n130 | 534.489 | 487.64 | | 566.54 | 0 | t_n130 | 572.171 | 563.62 | | 582.18 | 0 |
| t_n131 | 332.443 | 228.57 | | 429.15 | 0 | t_n131 | 507.857 | 435.36 | | 540.28 | 0 |
| t_n132 | 281.402 | 213.89 | | 351.7 | 0 | t_n132 | 363.643 | 163.27 | | 473.24 | 0 |
| t_n133 | 102.883 | 66.28 | | 149.85 | 0 | t_n133 | 150.452 | 50.94 | | 280.77 | 0 |
| t_n134 | 156.706 | 105.05 | | 217 | 0 | t_n134 | 262.718 | 98.67 | | 399.38 | 0 |
| T_n135 EUMETAZOA | 572.19 | 569.82 | | 573.07 | B(5.6100,5.7310,0.0250,1e-300) | T_n135 EUMETAZOA | 572.445 | 570.7 | | 573.08 | B(5.6100,5.7310,0.0250,1e-300) |
| t_n136 | 129.759 | 74.81 | | 201.71 | 0 | t_n136 | 176.552 | 65.85 | | 356.75 | 0 |
| t_n137 | 558.605 | 554.02 | | 562.85 | 0 | t_n137 | 557.065 | 554.03 | | 560.02 | 0 |
| T_n138  BILATERIA | 533.511 | 532.04 | | 537.29 | L(5.3200,0.1000,0.1000,1e-300) | T_n138  BILATERIA | 533.705 | 532.05 | | 538.14 | L(5.3200,0.1000,0.1000,1e-300) |
| T_n139  CNIDARIA | 533.606 | 531.85 | | 538.05 | L(5.3180,0.1000,0.1000,1e-300) | T_n139  CNIDARIA | 536.312 | 532.02 | | 542.72 | L(5.3180,0.1000,0.1000,1e-300) |

***Table S5*** MCMC diagnostics for time densities obtained with MCMCtree when fixing the tree topology used for BAMM analyses, both when the target distribution was set to be the prior (no data, label “CLK”) or the posterior (data were used, labels for relaxed-clock models are “AR” for autocorrelated rates and “IR” for independent rates sampled from a log-normal distribution)

| COI Topology used in BAMM | | | |
| --- | --- | --- | --- |
|  | **Prior-CLK** | **Autocorrelated- AR** | **Independent-IR** |
| tail-ESS times (median) | 49408 | 1198.5 | 1365.5 |
| tail-ESS times (min) | 49354 | 70.0 | 57.0 |
| tail-ESS times (max) | 49629 | 1464.0 | 2332.0 |
| bulk-ESS times (median) | 48943 | 1098.5 | 851 |
| bulk-ESS times (min) | 48880 | 42.0 | 44 |
| bulk-ESS times (max) | 49867 | 1538.0 | 2393 |
| Rhat min | 1.00008 | 0.9963348 | 0.9969007 |
| Rhat max | 1.000564 | 1.475644 | 3.844148 |
| Med. number of samples per chain | 20000 | 20000 | 20000 |
| Min. number of samples per chain | 20000 | 20000 | 20000 |
| Max. number of samples per chain | 20000 | 20000 | 20000 |
| Number of chains run | 64 | 64 | 64 |
| Number of chains kept after filters | 64 | 20 | 34 |
| Number of samples used to calculate tail-ESS and bulk-ESS | 100000 | 12512 | 11131 |
| Total number of samples kept *post-hoc* analyses | 100000 | 12512 | 11131 |

**Table S6**. Statistics obtained with BAMMTools after the BAMM on both divergence timetrees of Silicea with both molecular clocks (AR and IR).

| **Bamm stats** | **AR** | **IR** |
| --- | --- | --- |
| ESS postburn$N_shifts | 2126.471 | 13860.54 |
| ESS postburn$logLik | 1240.406 | 376.4842 |
| n.shifts | 2 | 2 |
| bulk-ESS times (median) | 1098.5 | 851 |
| bulk-ESS times (min) | 42.0 | 44 |

**Table S7** Species collected for the phylogenomic tree, time calibration and ASE analyses. Accession numbers will be added upon acceptance.

| S**pecies** | **Code** | **Order** | **Class** | **Accession number** |
| --- | --- | --- | --- | --- |
| *Leucetta giribeti* | LGIR | Clathrinida | Calcarea | This study |
| *Clathrina coriacea* | CCOR | Clathrinida | Calcarea | SRR3417192 |
| *Sycon ciliatum* | SCIL | Leucosolenida | Calcarea | Fortunato et al., 2012 |
| *Sycon coactum* | SCOA | Leucosolenida | Calcarea | [SRP012620](https://trace.ncbi.nlm.nih.gov/Traces?study=SRP012620) |
| *Agelas schmidtii* | ASCH | Agelasida | Demospongiae | This study |
| *Cymbaxinella damicornis* | CDAM | Agelasida | Demospongiae | [SRP224770](https://trace.ncbi.nlm.nih.gov/Traces?study=SRP224770) |
| *Cymbastela concentrica* | CCON | Bubarida | Demospongiae | [SRP061923](https://trace.ncbi.nlm.nih.gov/Traces?study=SRP061923) |
| *Phakellia robusta* | PROB | Bubarida | Demospongiae | [SRP271794](https://trace.ncbi.nlm.nih.gov/Traces?study=SRP271794) |
| *Phakellia ventilabrum* | PVEN | Bubarida | Demospongiae | [SRP268794](https://trace.ncbi.nlm.nih.gov/Traces?study=SRP268794) |
| *Chondrilla caribensis* | CCAR | Chondrillida | Demospongiae | [SRP012620](https://trace.ncbi.nlm.nih.gov/Traces?study=SRP012620) |
| *Cliona varians* | CVAR | Clionaida | Demospongiae | [SRP028615](https://trace.ncbi.nlm.nih.gov/Traces?study=SRP028615) |
| *Spheciospongia vesparium* | SVES | Clionaida | Demospongiae | SRR7702355 |
| *Dendrilla antarctica* | DANT | Dendroceratida | Demospongiae | [SRP192036](https://trace.ncbi.nlm.nih.gov/Traces?study=SRP192036) |
| *Ircinia fasciculata* | IFAS | Dictyoceratida | Demospongiae | [SRP012620](https://trace.ncbi.nlm.nih.gov/Traces?study=SRP012620) |
| *Vaceletia* sp. | VACSP | Dictyoceratida | Demospongiae | SRR4423080 |
| *Spongia officinalis* | SOFF | Dictyoceratida | Demospongiae | [SRP150632](https://trace.ncbi.nlm.nih.gov/Traces?study=SRP150632) |
| *Halisarca caerulea* | HCAE | Halisarcida | Demospongiae | [SRP098972](https://trace.ncbi.nlm.nih.gov/Traces?study=SRP098972) |
| *Halisarca dujardinii* | HDUJ | Halisarcida | Demospongiae | ERR1143554 |
| *Amphimedon queenslandica* | AQUE | Haplosclerida | Demospongiae |  |
| *Haliclona penicillata* | HPEN | Haplosclerida | Demospongiae | This study |
| *Petrosia ficiformis* | PFIC | Haplosclerida | Demospongiae | [SRP012620](https://trace.ncbi.nlm.nih.gov/Traces?study=SRP012620) |
| *Crella elegans* | CELE | Poecilosclerida | Demospongiae | [SRP012620](https://trace.ncbi.nlm.nih.gov/Traces?study=SRP012620) |
| *Iophon unicorne* | IUNI | Poecilosclerida | Demospongiae | This study |
| *Kirkpatrickia variolosa* | KVAR | Poecilosclerida | Demospongiae | [SRP056162](https://trace.ncbi.nlm.nih.gov/Traces?study=SRP056162) |
| *Latrunculia apicalis* | LAPI | Poecilosclerida | Demospongiae | [SRP056162](https://trace.ncbi.nlm.nih.gov/Traces?study=SRP056162) |
| *Mycale laevis* | MLAE | Poecilosclerida | Demospongiae | [SRP252526](https://trace.ncbi.nlm.nih.gov/Traces?study=SRP252526) |
| *Mycale tridens* | MTRI | Poecilosclerida | Demospongiae | [SRP252526](https://trace.ncbi.nlm.nih.gov/Traces?study=SRP252526) |
| *Mycale acerata* | MACE | Poecilosclerida | Demospongiae | [SRP252526](https://trace.ncbi.nlm.nih.gov/Traces?study=SRP252526) |
| *Mycale lingua* | MLIN | Poecilosclerida | Demospongiae | This study |
| *Mycale phyllophyla* | MPHY | Poecilosclerida | Demospongiae | [SRP051131](https://trace.ncbi.nlm.nih.gov/Traces?study=SRP051131) |
| *Isodyctia* sp. | ISOSP | Poecilosclerida | Demospongiae | [SRP120971](https://trace.ncbi.nlm.nih.gov/Traces?study=SRP120971) |
| *Crambe crambe* | CCRA | Poecilosclerida | Demospongiae | This study |
| *Phorbas areolatus* | PARE | Poecilosclerida | Demospongiae | [SRP252526](https://trace.ncbi.nlm.nih.gov/Traces?study=SRP252526) |
| *Chondrocladia robertballardi* | CROB | Poecilosclerida | Demospongiae | This study |
| *Tedania anhelans* | TANH | Poecilosclerida | Demospongiae | [SRP061923](https://trace.ncbi.nlm.nih.gov/Traces?study=SRP061923) |
| *Polymastia invaginata* | PINV | Polymastiida | Demospongiae | This study |
| *Scopalina ruetzleri* | SRUE | Scopalinida | Demospongiae | This study |
| *Scopalina* sp. | SCOPSP | Scopalinida | Demospongiae | [SRP061923](https://trace.ncbi.nlm.nih.gov/Traces?study=SRP061923) |
| *Spongilla lacustris* | SLAC | Spongillida | Demospongiae | [SRP012620](https://trace.ncbi.nlm.nih.gov/Traces?study=SRP012620) |
| *Ephydatia muelleri* | EMUE | Spongillida | Demospongiae | [SRP371964](https://trace.ncbi.nlm.nih.gov/Traces?study=SRP371964) |
| *Eunapius fragilis* | EFRA | Spongillida | Demospongiae | https://era.library.ualberta.ca/items/6139a88f-895d-44a7-bd0d-e22d455d2785 |
| *Lubomirskia baikalensis* | LBAI | Spongillida | Demospongiae | PRJNA431612 |
| *Lubomirskia abietina* | LABI | Spongillida | Demospongiae | PRJNA431612 |
| *Baikalospongia bacillifera* | BBAC | Spongillida | Demospongiae | PRJNA431612 |
| *Pseudospongosorites suberitoides* | PSUB | Suberitida | Demospongiae | [SRP037540](https://trace.ncbi.nlm.nih.gov/Traces?study=SRP037540) |
| *Prosuberites* sp. | PROSP | Suberitida | Demospongiae | This study |
| *Halichondria panicea* | HPAN | Suberitida | Demospongiae | [SRP100065](https://trace.ncbi.nlm.nih.gov/Traces?study=SRP100065) |
| *Geodia barretti* | GBAR | Tetractinellida | Demospongiae | [SRP246203](https://trace.ncbi.nlm.nih.gov/Traces?study=SRP246203) |
| *Geodia atlantica* | GATL | Tetractinellida | Demospongiae | [SRP246203](https://trace.ncbi.nlm.nih.gov/Traces?study=SRP246203) |
| *Geodia macandrewi* | GMAC | Tetractinellida | Demospongiae | [SRP246203](https://trace.ncbi.nlm.nih.gov/Traces?study=SRP246203) |
| *Geodia parva* | GPAR | Tetractinellida | Demospongiae | [SRP246203](https://trace.ncbi.nlm.nih.gov/Traces?study=SRP246203) |
| *Geodia phlegraei* | GPHL | Tetractinellida | Demospongiae | [SRP246203](https://trace.ncbi.nlm.nih.gov/Traces?study=SRP246203) |
| *Geodia hentscheli* | GHEN | Tetractinellida | Demospongiae | [SRP246203](https://trace.ncbi.nlm.nih.gov/Traces?study=SRP246203) |
| *Poecillastra compressa* | PCOM | Tetractinellida | Demospongiae | [SRP224770](https://trace.ncbi.nlm.nih.gov/Traces?study=SRP224770) |
| *Stelletta normani* | SNOR | Tetractinellida | Demospongiae | This study |
| *Stryphnus fortis* | SFOR | Tetractinellida | Demospongiae | [SRP224770](https://trace.ncbi.nlm.nih.gov/Traces?study=SRP224770) |
| *Craniella infrequens* | CINF | Tetractinellida | Demospongiae | This study |
| *Tethya wilhelma* | TWIL | Tethyida | Demospongiae | SRR4255675 |
| *Aplysina aerophoba* | AAER | Verongiida | Demospongiae | ERR2220853 |
| *Rossella fibulata* | RFIB | Lyssacinosida | Hexactinellida | [SRP056162](https://trace.ncbi.nlm.nih.gov/Traces?study=SRP056162) |
| *Sympagella nux* | SNUX | Lyssacinosida | Hexactinellida | [SRP056162](https://trace.ncbi.nlm.nih.gov/Traces?study=SRP056162) |
| *Vazella pourtalesii* | VPOU | Lyssacinosida | Hexactinellida | SRS6494416 |
| *Corticium candelabrum* | CCAN | Homosclerophorida | Homoscleromorpha | [SRP012620](https://trace.ncbi.nlm.nih.gov/Traces?study=SRP012620) |
| *Oscarella pearsei* | OPEA | Homosclerophorida | Homoscleromorpha | SRR1042040 |
| *Pleurobrachia bachei* | PBAC | Outgroup | Ctenophora | GCA_000695325.1 |
| *Bolinopsis ashley* | BASH | Outgroup | Ctenophora | SRR5892570 |
| *Nematostella vectensis* | NVEC | Outgroup | Cnidaria | ASM20922v1 |
| *Aurelia aurita* | AAUR | Outgroup | Cnidaria | SRR10240021 |
| *Lottia gigantea* | LGIG | Outgroup | Mollusca | GCF_000327385.1 |
| *Homo sapiens* | HSAP | Outgroup | Chordata | GCA_000306695.2 |

**Table S8** Convergence values for the two bayesian chains estimated with Tracecomp (Burnin = 1500), as implemented in Phylobayes

| **Name** | **Effective sample size** | **Real_diff** |
| --- | --- | --- |
| **Log-likelihood** | 61 | 0.15233 |
| **Tree length** | 709 | 0.0709294 |
| **Alpha parameter of gamma distribution** | 595 | 0.0110885 |
| **Number of modes** | 169 | 0.224488 |
| **State of entropy** | 316 | 0.099967 |
| **State alpha** | 1047 | 0.138845 |

**Table S9** Spicule types of the species included in the phylogenomic dataset, used to build the morphological matrix for the ASE. Spicule character information was taken from Systema Porifera.

| **Class/Order** | ***Species*** | **Megasclere** | **Microsclere** | **Mineralogy** |
| --- | --- | --- | --- | --- |
| **Agelasida** | *Agelas schmidtii* | stylotes | NO | Siliceous |
| **Agelasida** | *Cymbaxinella damicornis* | oxeas | NO | Siliceous |
| **Bubarida** | *Cymbastela concentrica* | oxeas | NO | Siliceous |
| **Bubarida** | *Phakellia robusta* | oxeas | NO | Siliceous |
| **Bubarida** | *Phakellia ventilabrum* | oxeas | NO | Siliceous |
| **Chondrillida** | *Chondrilla caribensis* | NO | spherasters, oxyspherasters (microspination on the tips) | Siliceous |
| **Clionaida** | *Cliona varians* | tylostyles | spirasters | Siliceous |
| **Clionaida** | *Spheciospongia vesparium* | tylostyles | spirasters | Siliceous |
| **Dendroceratida** | *Dendrilla antarctica* | NO | NO | Absent |
| **Dictyoceratida** | *Ircinia fasciculata* | NO | NO | Absent |
| **Dictyoceratida** | *Spongia officinalis* | NO | NO | Absent |
| **Dictyoceratida** | *Vaceletia* sp. | NO | NO | Aragonite |
| **Halisarcida** | *Halisarca caerulea* | NO | NO | Absent |
| **Halisarcida** | *Halisarca dujardinii* | NO | NO | Absent |
| **Haplosclerida** | *Amphimedon queenslandica* | oxeas | NO | Siliceous |
| **Haplosclerida** | *Haliclona penicillata* | oxeas | NO | Siliceous |
| **Haplosclerida** | *Petrosia ficiformis* | oxeas | NO | Siliceous |
| **Poecilosclerida** | *Chondrocladia robertballardi* | Mycalostyles | isochaele | Siliceous |
| **Poecilosclerida** | *Crambe crambe* | Subtylostyles, styles | unguiferate anchorate chelae (28–(38.3)–43m) astrose desmoid spicules rounded knobbed ends, cladome 38–(68.5)–99, rays 10–(35.7)–477 (10.6)–14m | Siliceous |
| **Poecilosclerida** | *Crella elegans* | Tornotes, oxeote; acanthostyles, acanthoxeas | NO | Siliceous |
| **Poecilosclerida** | *Iophon unicorne* | Choanosomal oxeote styles, Ectosomal strongyles | Bipocoelles, Anisochelae | Siliceous |
| **Poecilosclerida** | *Isodictya* sp. | oxeas | isochelae | Siliceous |
| **Poecilosclerida** | *Kirkpatrickia variolosa* | styles, strongyles | NO | Siliceous |
| **Poecilosclerida** | *Latrunculia apicalis* | Smooth styles | aciculodiscorhabds | Siliceous |
| **Poecilosclerida** | *Mycale (Oxymycale) acerata* | oxeas | anisochelae, trichodragmas | Siliceous |
| **Poecilosclerida** | *Mycale (Mycale) laevis* | oxeas | anisochelae, trichites, anisanchorae | Siliceous |
| **Poecilosclerida** | *Mycale (Mycale) lingua* | styles/mycalostyles | Anisochelae, sigmas, raphide | Siliceous |
| **Poecilosclerida** | *Mycale (Carmia) phyllophila* | tylostyle/subtylostyles | Anisochelae, sigmas | Siliceous |
| **Poecilosclerida** | *Mycale (Mycale) tridens* | Mycalostyles | Chaele | Siliceous |
| **Poecilosclerida** | *Phorbas areolatus* (antarctica) | acanthostyles, amphioxeas | isochelae | Siliceous |
| **Poecilosclerida** | *Tedania (Tedania) anhelans* | Styles, tylostyles | onychaetes | Siliceous |
| **Polymastiida** | *Polymastia invaginata* | Styles, tylostyles | NO | Siliceous |
| **Scopalinida** | *Scopalina ruetzleri* | styles with oxeote modifications | NO | Siliceous |
| **Scopalinida** | *Scopalina* sp. | styles | NO | Siliceous |
| **Spongillida** | *Baikalospongia bacillifera* | slightly curved amphistrongyla, spines near their ends, rarely oxeas (26227m), | NO | Siliceous |
| **Spongillida** | *Ephydatia fluviatilis* | acanthoxeas, slightly curved or rarely straight oxeas, from smooth to microspined (210–4006–19m). | gemmoscleres Microscleres absent | Siliceous |
| **Spongillida** | *Eunapius fragilis (emue)* | oxeas | gemmoscleres (spiny amphistrongyla) | Siliceous |
| **Spongillida** | *Lubomirskia abietina* | curved amphioxea, fusiform, spines near their tips | No mention | Siliceous |
| **Spongillida** | *Lubomirskia baikalensis* | oxeas, acanthoxeas, slightly curved amphioxea, fusiform, many spines along whole length | Absent | Siliceous |
| **Spongillida** | *Spongilla lacustris* | oxeas tips gently to sharply pointed, slightly spined if associated to gemmules, acanthoxeas | fusiform oxeas (25–1782–8m) with dense spines regularly distributed along length and microspinosity on spines , asterose shape. Gemmules two types in the same specimen, naked without gemmuloscleres and armoured with gemmuloscleres | Siliceous |
| **Suberitida** | *Halichondria panicea* | oxeas | NO | Siliceous |
| **Suberitida** | *Pseudospongosorites suberitoides* | smooth sharp pointed oxeas, occasionally centrotylote and bent, occasionally stylote or strongylote, probably divisible in two overlapping size categories, 176–2952–12m and 125–1852–10m. | No mention | Siliceous |
| **Suberitida** | *Suberites domuncula* | tylostyles, slightly subterminal (drop-shaped) but mostly well-formed tyle, except for rare annular swellings in the neck region. Small surface tylostyles, 100–3504–8m, larger choanosomal tylostyles 250–4805–8m, | No mention | Siliceous |
| **Tethyida** | *Tethya wilhelma* | strongyloxeas, anisostrongyles, styles | spherasters, tylasters, strongylasters | Siliceous |
| **Tetractinellida** | *Craniella infrequens* | anatriaenes, protriaenes, oxeas | NO | Siliceous |
| **Tetractinellida** | *Geodia barretti* | oxeas, anatriaenes, dichotriaene | sterrasters, oxyaster, strongylaster, microxea | Siliceous |
| **Tetractinellida** | *Geodia atlantica* | oxeas, dichotriaenes, orthotriaenes | sterrasters, spheroxyasters, oxyasters | Siliceous |
| **Tetractinellida** | *Geodia hentscheli* | oxeas, dichotriaenes, mesoprotriaenes | sterrasters, oxyaster, strongylaster, microxea | Siliceous |
| **Tetractinellida** | *Geodia macandrewii* | oxeas, anatriaenes, orthotriaenes, promesotriaene | sterrasters, spheroxyasters, oxyasters | Siliceous |
| **Tetractinellida** | *Geodia parva* | oxeas, anatriaenes, orthotriaenes, Meso/protriaenes | sterrasters, spherasters, oxyasters | Siliceous |
| **Tetractinellida** | *Geodia phlegraei* | oxeas, anatriaenes, orthotriaenes, Meso/protriaenes | sterrasters, spherasters, oxyasters | Siliceous |
| **Tetractinellida** | *Poecillastra compressa* | oxeas, short-shated triaenes | microxeas, spiraster to metaster, plesiaster | Siliceous |
| **Tetractinellida** | *Stelletta normanii* | oxeas, anatriaenes, dichotriaenes, protriaenes | oxyaster, strongylaster, trichodragma | Siliceous |
| **Tetractinellida** | *Stryphnus fortis* | oxeas, dichotriaenes, plagiotriaenes, anatriaenes, mesoanatriaenes | oxyasters, sanidasters to amphisanidasters | Siliceous |
| **Verongiida** | *Aplysina aerophoba* | NO | NO | Absent |
| **Homosclerophorida** | *Corticium candelabrum* | Chaltrops | Microchaltrops | Siliceous |
| **Homosclerophorida** | *Oscarella pearsei* | NO | NO | Absent |
| **Lyssacinosida** | *Rossella fibulata* | heterodiactin, pentactin, diactins | oxyhexasters | Siliceous |
| **Lyssacinosida** | *Sympagella nux* | pinular hexactin, pentactin, diactins | Discohexasters | Siliceous |
| **Lyssacinosida** | *Vazella pourtalesii* | stauractin, pentactin, diactins | microdiscohexasters, Hemihexasters | Siliceous |
| **Calcarea** | *Clathrina coriacea* | triactines equiangular and equiradiate |  | Calcite |
| **Calcarea** | *Leucetta giribeti* | triactines and tetractines |  | Calcite |
| **Calcarea** | *Sycon ciliatum* | triactines |  | Calcite |
| **Calcarea** | *Sycon coactum* | triactines and tetractines |  | Calcite |

**Table S10** ST calibrations applied to the COI topology used for the diversification rates analyses in BAMM and MEDUSA

| **Name of clades corresponding to nodes** | **node** | **Prior** |
| --- | --- | --- |
| **Silicea** | 808 | ST(5.8300,0.0590,0.1120,109.1240) |
| **Demospongiae** | 809 | ST(5.4510,0.0700,0.9220,209.2980) |
| **Haplosclerida + + Heteroscleromorpha** | 810 | ST(5.1500,0.0190,394.1320,3.5100) |
| **After split with Haplosclerida** | 811 | ST(4.5960,0.2010,-1.1000,404.5290) |
| **After split with Scopalinida +Spongillida** | 812 | ST(4.0910,0.1770,-0.3850,246.6070) |
| **After split with Bubarida** | 813 | ST(3.6390,0.1960,0.7740,151.2750) |
| **After split with Agelasida** | 815 | ST(3.2620,0.2050,1.1200,431.6690) |
| **Poecilosclerida** | 928 | ST(2.6670,0.2050,1.5490,477.3950) |
| **Bubarida** | 1143 | ST(0.6040,0.1810,2.1930,136.9570) |
| **Tetractinellida** | 1179 | ST(2.4770,0.2300,1.4590,701.2120) |
| **Spongillida + Scopalinida** | 1353 | ST(4.0020,0.3260,-1.4770,1215.9540) |
| **Haplosclerida** | 1392 | ST(3.5230,0.4820,-0.9020,983.7760) |
| **Hexactinellida** | 1563 | ST(4.4500,0.0330,1588.4600,3.5420) |

**Table S11** Sample frequencies used for the BAMM and MEDUSA analysis

| **Clade** | **Species Sampled** | **Num. species accepted** | **Sampling frquency** |
| --- | --- | --- | --- |
| **Silicea** | 807 | 8649 | 0.09330558 |
| **Hexactinellida** | 52 | 701 | 0.07417974 |
| **Agelasida** | 26 | 73 | 0.35616438 |
| **Tetractinellida** | 175 | 1175 | 0.14893617 |
| **Haplosclerida** | 66 | 1147 | 0.05754141 |
| **Poecilosclerida** | 165 | 2448 | 0.06740196 |
| **Polymastiida** | 33 | 132 | 0.25 |
| **Axinellida** | 25 | 534 | 0.04681648 |
| **Bubarida** | 12 | 153 | 0.07843137 |
| **Spongillida** | 33 | 283 | 0.11660777 |
| **Tethyida** | 27 | 212 | 0.12735849 |
| **Biemnida** | 10 | 106 | 0.09433962 |
| **Sphaerocladina** | 4 | 5 | 0.8 |
| **Dictyoceratida** | 51 | 588 | 0.08673469 |
| **Dendorceratida** | 17 | 70 | 0.24285714 |
| **Verongiida** | 31 | 95 | 0.32631579 |
| **Scopalinida** | 3 | 39 | 0.07692308 |
| **Suberitida** | 40 | 519 | 0.07707129 |
| **Chondrosida** | 2 | 11 | 0.18181818 |
| **Chondrillida** | 6 | 42 | 0.14285714 |
| **Clionaida** | 25 | 213 | 0.11737089 |
| **Desmacellida** | 3 | 37 | 0.08108108 |

Data S1. (separate file)

Table with all the ASE results for each topology analysed.
